# OfUSA: OpenfUS Analyzer, a versatile open-source framework for the analysis and visualization of functional ultrasound imaging data across animal models

**DOI:** 10.1101/2025.09.16.676515

**Authors:** Yun-An Huang, Théo Lambert, Damon Verbeyst, Nora Eilis Fitzgerald, Micheline Grillet, Clément Brunner, Gabriel Montaldo, Wim Vanduffel, Alan Urban

**Author notes:** Corresponding author: Alan Urban.

## Abstract

Functional ultrasound (fUS) imaging is rapidly gaining interest for its unprecedented ability to study large-scale brain dynamics, yet its adoption and broader dissemination have been hindered by the lack of standardized tools and methodologies to analyze and interpret its rich datasets. We present OpenfUS Analyzer (OfUSA), a companion software suite designed to help researchers quickly engage with fUS data and perform the full range of analyses needed to generate publication-ready results and figures without relying on additional software. OfUSA offers a versatile and modular architecture including preprocessing, recording quality assessment, signal dynamics exploration, statistical analysis and visualization. These functions are separated yet easily combined into analytic pipelines through a programming-free graphical interface. The framework can be applied across species and experimental contexts, either by registering data to anatomical atlases, as shown here for the mouse brain, or by analyzing data without atlas constraints, as illustrated in a primate dataset. This flexibility, together with its comprehensive functionality, makes OfUSA a practical solution for standardized and reproducible analysis of fUS data in both preclinical and translational research. Using OfUSA, we demonstrate the capacity to detect stimulus-evoked responses with high sensitivity, identify their spatial localization within brain networks, and quantify both their extent and temporal dynamics. These results highlight the software’s ability to capture robust activation patterns and provide detailed insights into brain function, thereby accelerating the use of fUS as a powerful tool for systems neuroscience.

**Highlights:** We present OpenfUS Analyzer (OfUSA), a novel software platform for the complete analysis of functional ultrasound (fUS) datasets. OfUSA combines a user-friendly graphical interface with a standardized, flexible workflow and powerful visualization tools, making it an ideal solution for fUS researchers at all experience levels. The software’s utility is first demonstrated through the analysis of rodent fUS data using a standardized atlas, while its versatility is further emphasized by the successful analysis of a primate fUS dataset without a template, thereby illustrating its adaptability to non-standard experimental conditions.

## Introduction

Imaging large-scale and in-depth brain activity in behaving rodents remains a significant challenge in neuroscience. Functional ultrasound imaging (fUS) overcomes many of these limitations by enabling measurements of hemodynamic responses across several cubic centimeters of brain tissue, with both high spatial and temporal resolution (around 100 μm^3^ and ∼100 ms, respectively) in awake rodents (Brunner et al., 2020; Montaldo et al., 2022). In comparison, optical imaging and electrophysiological recordings are restricted to localized regions, while fMRI is hindered by lower spatiotemporal resolution and the frequent requirement for anesthesia. Although awake-rodent fMRI has been demonstrated (Brydges et al., 2013; King et al., 2005; Zhang et al., 2010), it remains technically demanding and uncommon.

With the unique ability of fUS to capture large-scale activity across the awake brain, the increasing maturity of the technology, and its growing accessibility to the neuroscience community, there is a rising need for standardized and user-friendly tools to analyze fUS datasets. This need is driven by two main factors. First, the rapid expansion of fUS applications is generating a steadily increasing volume of data. fUS has already been applied to investigate the visual pathway (Brunner et al., 2020; Gesnik et al., 2017; É. Macé et al., 2018), tactile processing in whiskers (E. Macé et al., 2011), olfactory perception (B. F. Osmanski et al., 2014), resting-state dynamics (B.-F. Osmanski et al., 2014), stroke (Brunner et al., 2024, 2017; Hingot et al., 2020), pain (Claron et al., 2021), and drug effects (Rabut et al., 2020; Vidal, Droguerre, Valdebenito, et al., 2020; Vidal, Droguerre, Venet, et al., 2020). In addition, fUS is highly compatible with other techniques, such as optogenetic stimulation (Brunner et al., 2020; Rungta et al., 2017; Sans-Dublanc et al., 2021) and electrophysiological recordings (Lambert, Niknejad, et al., 2025; E. Macé et al., 2011; Nunez-Elizalde et al., 2022; Sans-Dublanc et al., 2021), thereby enabling multimodal investigations of brain function. This breadth and diversity of applications require analytical pipelines that are not only standardized and reproducible but also flexible and adaptable while maintaining robustness.

Second, there is a notable absence of open-source analysis software specifically tailored to the needs of fUS researchers. Although commercial scanners generally provide embedded software, these solutions are proprietary, limiting accessibility, transparency, and customization for the broader neuroscience community (Bertolo et al., 2021). Our OpenfUS initiative has recently introduced PyfUS, a collection of scripts and analysis tools designed to process fUS data at the voxel level (Lambert, Brunner, et al., 2025). While it provides valuable functionalities such as correlation and clustering analyses, its effective use requires users to set up a dedicated computing environment with the appropriate libraries and packages. As a result, adapting and operating PyfUS still demands considerable programming expertise, which restricts its accessibility for researchers without advanced computational skills. Existing open-source packages developed for fMRI, EEG, or optical imaging analysis also fail to meet the specific requirements of fUS. Despite sharing certain statistical methods, the preprocessing and visualization workflows differ substantially. For instance, fUS data lacks standardized quality control procedures, and many preprocessing steps used in fMRI are unnecessary for ultrasound signals. Similarly, registration methods designed for T1-, T2-, or EPI-based rodent fMRI cannot be directly applied to Doppler images because of their distinct contrast properties (Brunner et al., 2021). Moreover, fUS researchers focus not only on activation maps but also on the temporal dynamics of hemodynamic responses and the visualization of barcode-like spatiotemporal patterns, which are less emphasized in fMRI workflows (Brunner et al., 2021, 2023; Lambert, Brunner, et al., 2025; É. Macé et al., 2018). Altogether, these limitations highlight the urgent need for an open-access, unified, and user-friendly framework that combines standardization with flexibility and is explicitly designed for the analysis of fUS data.

In this work, we introduce the OpenfUS Analyzer (OfUSA), an open-access software platform specifically developed to support researchers in preprocessing, statistical analyses, and visualizing fUS data. Expanding on scripts from our previous work (Brunner et al., 2021), OfUSA significantly enhances usability and scalability, enabling the processing of large datasets across multiple animals and experimental sessions. The software was designed with six core objectives that address critical needs of the neuroscience community: 1) streamline fUS analysis workflows to lower technical barriers for researchers; 2) establish standardized and reproducible pipelines that ensure methodological rigor and comparability across studies; 3) maintain flexibility to support diverse experimental paradigms and species; 4) guarantee interoperability with established third-party software environments; 5) provide an intuitive, user-friendly graphical interface that makes advanced analyses accessible without programming expertise; and 6) integrate a comprehensive suite of visualization tools to enable in-depth exploration and interpretation of brain activity.

To demonstrate its versatility across species, we illustrate the software’s application using datasets from rodents and non-human primates. Rodents were chosen as the primary example because they are the most widely used model in neuroscience, allowing readers to directly relate the workflow to their own research. In this case, the analysis is supported by a standardized template, which provides additional anatomical and vascular information to aid in data interpretation. As a complementary example, we present a dataset from non-human primates, highlighting one of the key opportunities of fUS: its application to gyrencephalic brains that can serve as a bridge toward human translation. In this scenario, no standardized template is available and only part of the brain is accessible, thereby demonstrating how the software can accommodate conditions typical of larger-brain studies. Together, these examples showcase both the flexibility of the analysis pipeline and the depth of spatial and temporal information that can be extracted.

To promote broader adoption and standardization of fUS, the software will be integrated into a web-based platform designed to serve as a global hub for the fUS community. This platform will streamline the entire workflow, from experimental design to advanced data analysis, while fostering collaboration, reproducibility, and innovation in brain research (**Figure 1a**).

**Figure 1.**
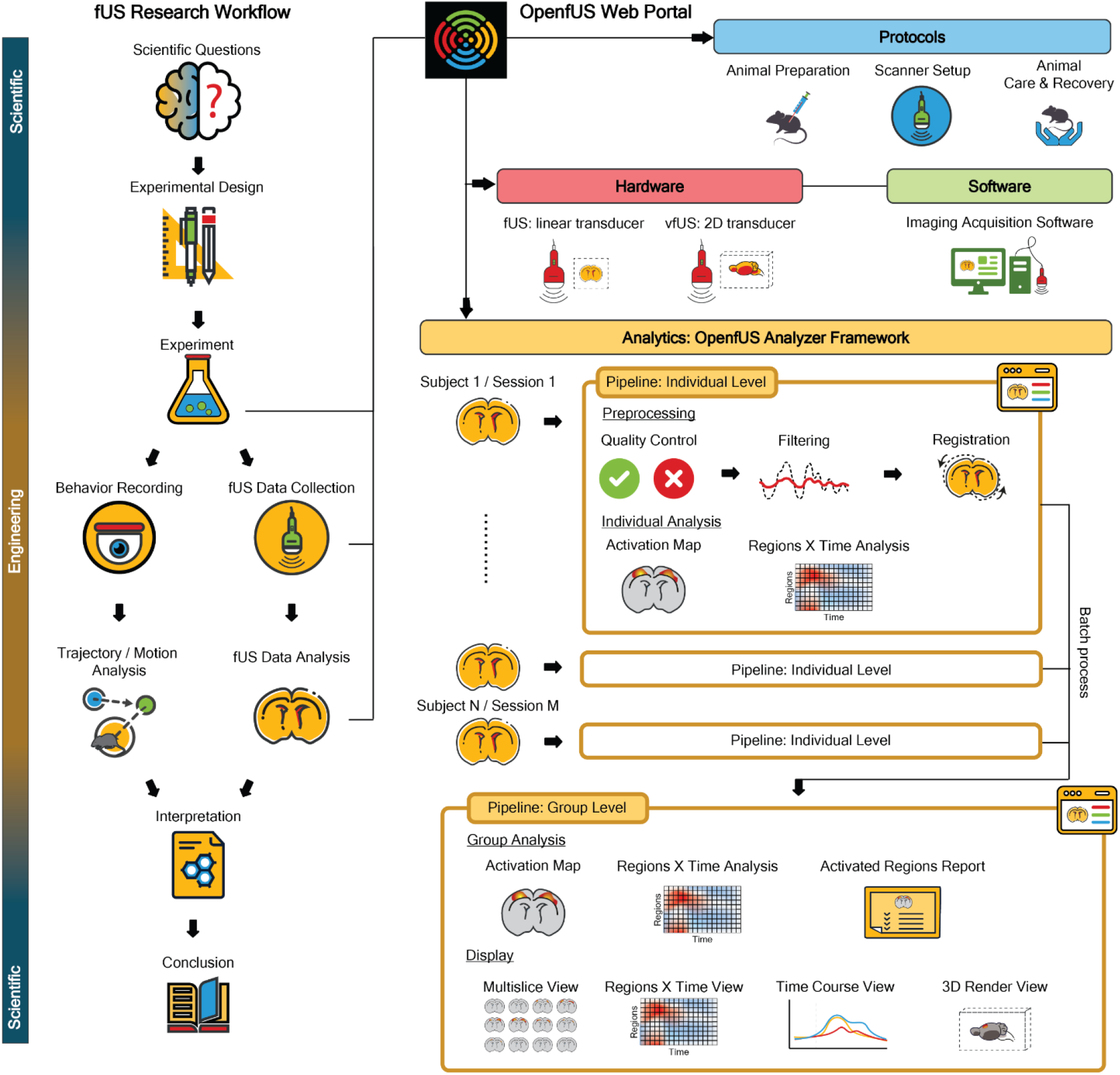
OpenfUS Web Portal and OfUSA Framework. This figure schematizes the workflow for functional ultrasound (fUS) research developed within the OpenfUS initiative. The left panel depicts the progression from scientific questions to final conclusions, with the initial and terminal phases relying on scientific expertise, and the intermediate stages requiring engineering expertise and substantial development efforts. The right panel displays the OpenfUS toolkit, which provides hardware specifications, experimental protocols, acquisition software, and analysis software to address the technical requirements of fUS research. Within this framework, OfUSA is introduced as a central software component, enabling standardized and user-friendly analysis and visualization of fUS data in rodent models. The standard workflow (bottom right) includes preprocessing, individual-level and group-level analyses, and visualization steps.

### OfUSA Overview

OfUSA is designed to analyze fUS datasets acquired from rodents or other animal models, capturing three spatial dimensions together with the temporal dynamics of cerebral blood flow. In practice, fUS data can be generated using different hardware configurations combining the transceiver, the ultrasound transducer, and the computing workstation. For clarity, this manuscript focuses on the most widely used setups: either a linear transducer with 128 channels or a matrix array with 1024 channels, enabling the acquisition of data restricted to a specific brain section (coronal or sagittal) or extending over a large portion of the rodent’s brain. Each voxel in an fUS dataset encodes a signal proportional to cerebral blood volume (CBV), which has been shown to correlate closely with underlying neuronal spiking activity (Lambert, Niknejad, et al., 2025; É. Macé et al., 2018; Montaldo et al., 2022; Nunez-Elizalde et al., 2022), thereby providing a robust link between hemodynamic signals and neural function.

The analysis aims to unveil the spatial information and the temporal information of hemodynamic response evoked by the external stimuli (task). The spatial information refers to the distribution of the activated regions that are induced by external stimuli. We can examine the activated region by estimating the signal change between the baseline period and the stimulated period in each brain voxel. By applying a threshold, we can generate an activation map that highlights the regions with significant signal change. The temporal information refers to the characteristic of the hemodynamic response (changes of cerebral blood volume) time course in the regions of interest (ROI), which includes the shape of time course, rising time and the peak amplitude. We can extract time courses by averaging time courses across all voxels in ROIs. The default ROIs are based on Allen Brain Atlas for rodent or researchers can customize the ROIs in the software. Although the spatial information and temporal information are independent aspects, in practice, we can examine the activation map first to identify the stimuli-related ROIs, then we delve into the temporal information of hemodynamic response in those specific ROIs.

To facilitate the analysis of spatial and temporal information without imposing a heavy technical burden, OfUSA was developed in MATLAB®, a programming environment widely used in academia, with licenses often provided at reduced cost or free of charge through institutional agreements. The software integrates a graphical user interface (GUI) that streamlines the entire analysis workflow (**Figure 2**). The interface is organized into three main components. The first is the project management panel, which handles the basic settings of the project, the organization of data and process libraries, and the construction of analysis pipelines. A pipeline corresponds to a predefined sequence of processes that can be saved and reused, and several built-in pipeline templates are provided to help researchers quickly initiate their analyses. The second component is the process management panel, where users can edit process-specific parameters and assign datasets to be processed. Depending on the parameter type, inputs can be entered via text boxes, multi-choice menus, or dedicated interfaces for selecting files and directories. The third component is the visualization panel, which allows researchers to monitor logs and flexibly display results. Users can adjust visualization settings, such as color bar range, thresholds, and colormaps, enabling tailored exploration and clear presentation of figures.

**Figure 2.**
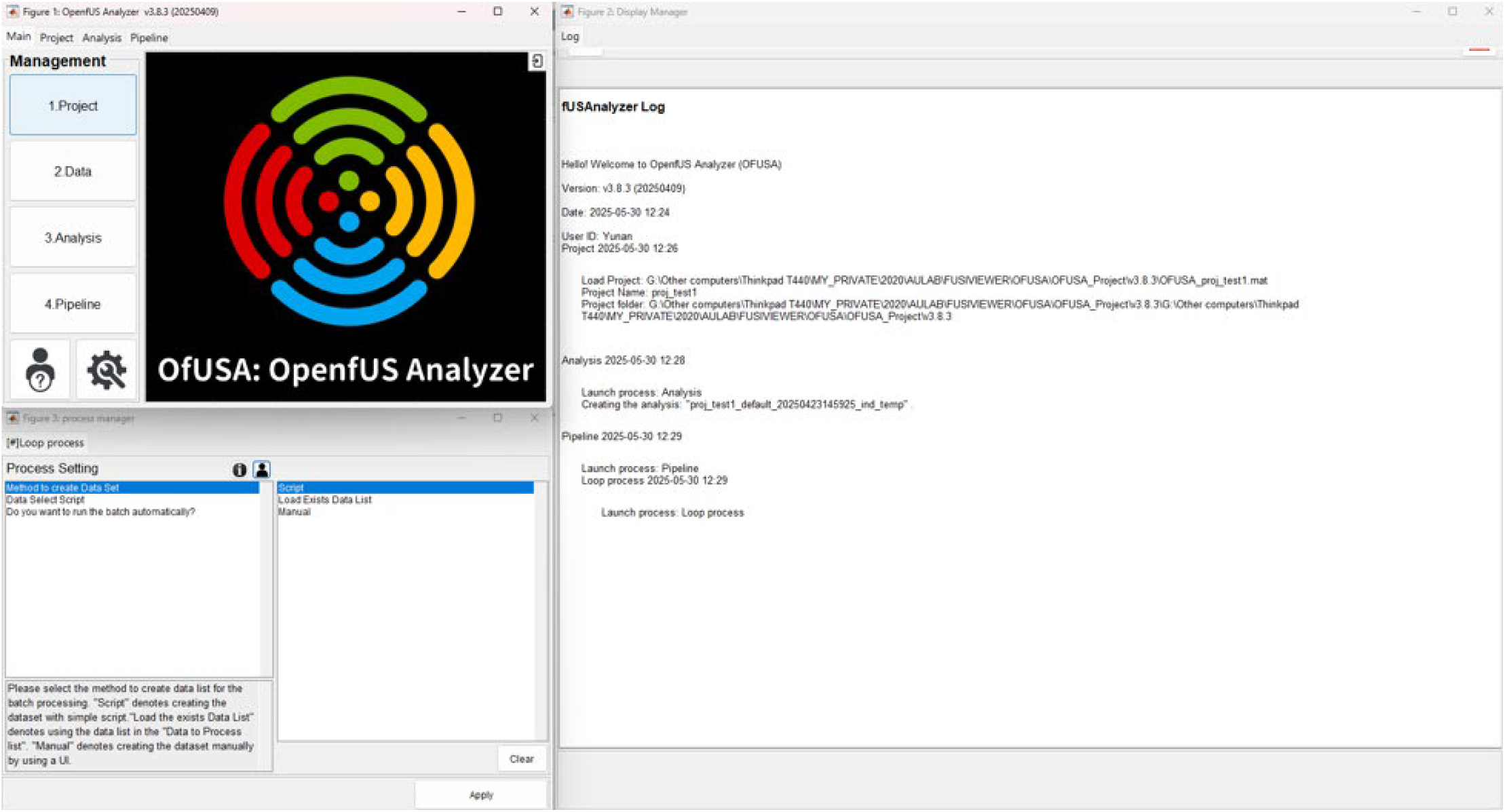
Graphical User Interface of OfUSA. The interface comprises three interactive panels for workflow management, process management, and data visualization. The top-left panel provides project management functions, including project setup, data loading, analysis creation, and the application of pipeline templates. The bottom-left panel manages process management, where analytical parameters are listed in a table and can be modified through corresponding input fields, selection menus, or file and directory dialogues. The right panel serves as the visualization area for displaying logs and data.

To guarantee seamless compatibility with third-party software, OfUSA stores all processed data in the NIFTI+JSON format, a standard widely adopted in the neuroscience community. NIFTI was specifically developed for storing 3D and 4D medical images or matrices, providing an efficient and structured way to handle large volumetric datasets. Complementing this, JSON is a lightweight and human-readable format for metadata, organized as attribute–value pairs. Its simplicity ensures that metadata can be accessed and edited with any text editor, making it highly convenient for data sharing, reproducibility, and integration into diverse workflows. In addition, raw fUS data are organized according to the Brain Imaging Data Structure (BIDS), a community-driven standard for structuring neuroimaging datasets. The use of BIDS not only promotes transparency and reproducibility but also facilitates interoperability across research groups by enabling straightforward data exchange and large-scale collaborative analyses. The full list of metadata stored in the JSON files is detailed in **Table S1**.

### Individual and Group analysis

The analysis workflow can be divided into two main stages, as illustrated in **Figure 1**. In the first stage, the fUS data for each trial is preprocessed independently to account for variability in system noise, physiological state, and animal movement. Because these sources of variability differ across trials, quality must be assessed and noise removed on a trial-by-trial basis (or at the session level when data are acquired with a linear transducer). This stage, referred to as the individual-level stage, includes three preprocessing steps: quality control, filtering, and registration. Quality control and filtering reduce the influence of artifacts generated by either the animal or the acquisition system, while registration aligns the fUS data to a reference atlas, such as the Allen Brain Atlas, ensuring consistent anatomical localization across sessions and subjects. Following preprocessing, individual-level analyses are conducted to generate activation maps and extract time courses from regions of interest (ROIs) for each trial or session. To increase efficiency, these procedures are implemented through batch processing, allowing multiple sessions or subjects to be analyzed simultaneously.

The second stage investigates group-level effects by combining data across animals, sessions, and trials. At this level, statistical analyses are performed to characterize both fixed effects, representing stimulus-driven responses within the measured sample, and mixed effects, which generalize responses by accounting for inter-animal and inter-session variability. Group-level outputs include activation maps and ROI time courses derived from statistical approaches such as averaging and t-tests. Visualization tools integrated into the software further support interpretation, providing multislice displays, region-by-time plots, time course visualizations, and 3D renderings.

**Figure 3** illustrates the analytical workflow from the generation of individual activation maps and ROI time courses to their aggregation at the group level. Additional methodological details, including mathematical formulations used in this standardized pipeline, are provided in **Appendix I, Section A**.

**Figure 3.**
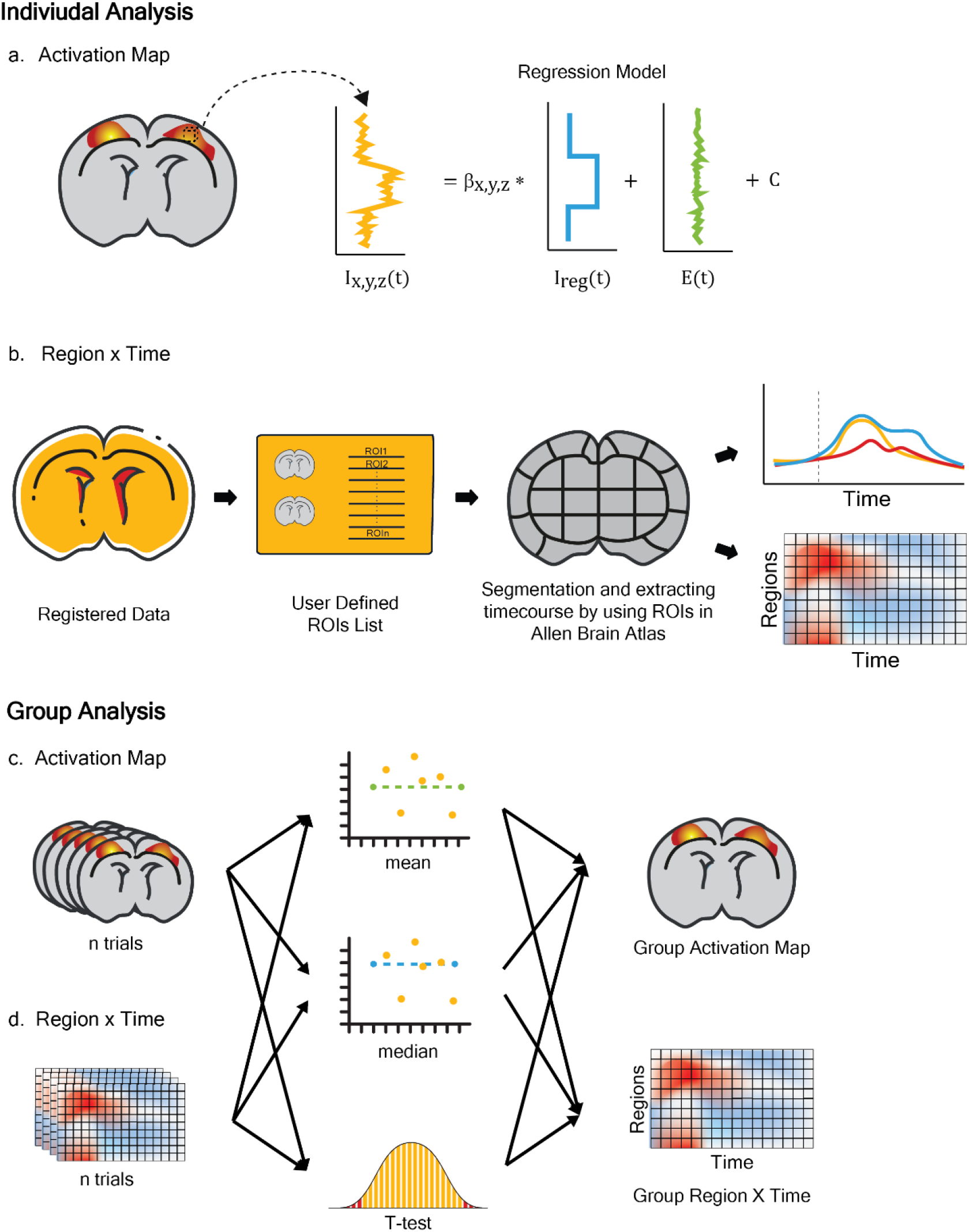
Concept of individual- and group-level analyses. Each dataset provide two primary outputs: activation maps and regional time courses. Activation maps represent the spatial distribution of signal changes associated with external stimuli, whereas regional time courses describe the temporal profile of cerebral blood flow changes within defined regions of interest. The upper row illustrates the generation of (a) activation maps using regression analysis (general linear model) and (b) regional time courses obtained by averaging voxel signals within each region for individual trials. The lower row illustrates the derivation of (c) activation maps and (d) regional time courses at the group level. Group-level analyses can be performed using mean or median averaging to assess fixed effects, or by applying a t-test to evaluate mixed effects while accounting for variability across animals and sessions.

## Application Example 1: Large-Scale Mapping of Evoked Brain Activity in Mice

### 1.1 Experimental setup

This multi-sensory experiment involved a wild-type mouse across eight imaging sessions. During each session, the mice were presented with two types of 5-second stimuli in a randomized order: a visual checkerboard and a tactile comb stimulation to the right whisker pad. Each stimulus condition was repeated 20 times within a 30-second trial structure that included a baseline (10s / 20 frames), stimulation (5s / 10 frames), and recovery period (15s / 30 frames) . A high-quality anatomical image (refers to a high resolution vascular images) was also acquired in each session for registration. **Figure 4a-b** depicts the mouse experimental setup and the stimulus. Further details on the experimental setup are provided in **Appendix II, Section A**.

**Figure 4.**
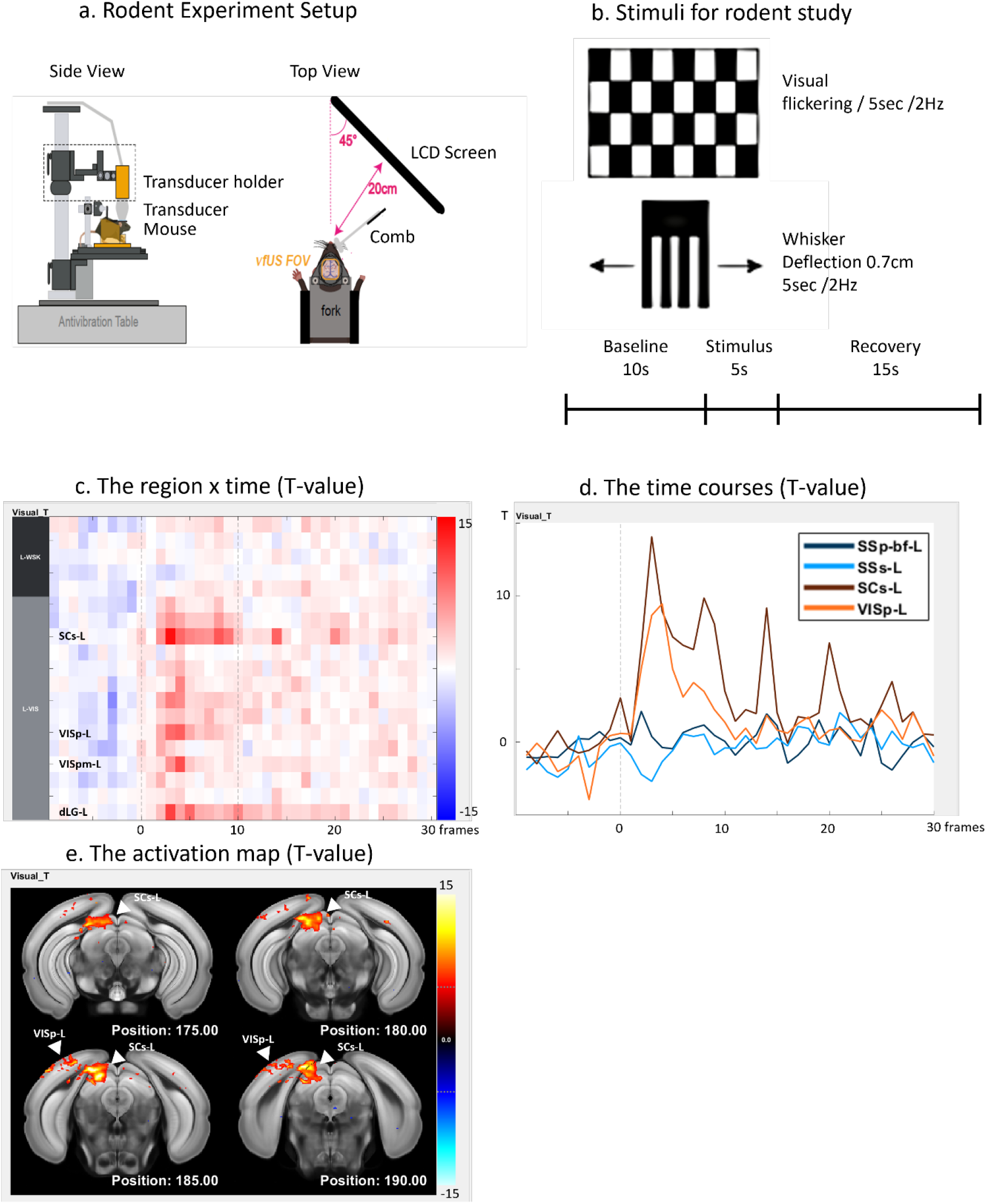
Experimental setups and results for the rodent study. Panel (a) shows the experimental setup for the sensory-evoked stimulation, with visual stimulation delivered through an LCD monitor positioned 20 cm in the right side of the animal and tactile stimulation applied to the right whiskers using a custom made comb. Panel (b) specifies the stimuli: a 2 Hz flickering checkerboard and a 0.7 cm comb deflection, each lasting 5 seconds, with before a baseline of 10s and after a recovery period of 15s. Panel (c) presents region-by-time plots of time courses across multiple brain regions. Panel (d) displays representative time courses for visual regions (VISp-L, SCs-L, orange) and somatosensory regions (SSp-bfd-L, SSs-L, blue). Panel (e) shows the group-level activation maps. Reported activations are statistically significant according to a one-sample t-test (T > 5.4, p < 0.001 uncorrected, df = 7, two-tailed).

### 1.2 Analysis Workflow

For the analysis of multi-sensory data, we utilized the standardized workflow provided by OfUSA, which is structured into two main stages: the individual-level stage and the group-level stage. The analysis began at the individual-level stage (**Figure S1a**), where data was processed on a trial-by-trial basis to obtain an average effect for each session. Subsequently, the group-level stage examined the random effects across these sessions to account for inter-session variability and produce corresponding statistical inferences (**Figure S1b**). This two-stage procedure effectively constitutes a mixed-effects model, allowing for the study of generalizable effects by incorporating session-to-session variability.

#### 1.2.1 Realignment

Prior to this two-stage analysis, all data were registered to the Allen Brain Atlas to ensure spatial alignment across sessions. We performed this alignment using the “registration ABA” process within the software. This process employed high-quality anatomical images to estimate the transformation matrix required to map each individual’s native space to the common Allen Brain Atlas space. **Figure S1c** illustrates the manual registration interface used for this purpose.

#### 1.2.2 Individual Level Analysis

**Figure S1a** presents the individual-level analysis pipeline, which is organized into four main components: initiation, preprocessing, temporal information analysis, and spatial information analysis.

The initiation component consists of three processes: “Loop Pipeline,” “Set Parameters,” and “Regressor.” The “Loop Pipeline” process defines the data units for each iteration of the batch process (e.g., subject, session, stimulus). Unlike in rodent studies, image data and regressors are selected within each loop. When multiple units, such as several subjects or sessions, are specified, the pipeline generates a corresponding number of independent processing loops (**Figure S1d**). The “Set Parameters” process specifies the regions of interest (ROIs) derived from the Allen Brain Atlas and assigns names to the experimental stimuli. In this study, 82 cortical and subcortical regions associated with multisensory processing were used, which, when divided by hemisphere, yielded 164 ROIs (**Table S2**). The “Regressor” process defines the experimental paradigm, including the onset of the stimulus (frame 20), duration (10 frames), and the expected shape of the time course (**Figure S1e**).

The preprocessing component includes quality control and filtering. During quality control, each trial is validated against thresholds to ensure robustness against motion artifacts and global noise while meeting minimal Signal-to-Noise Ratio (SNR) and Contrast-to-Noise Ratio (CNR) requirements. The thresholds applied were: TH_BurstError = 200%, TH_noisyvoxel = 50%, TH_SNR = 5, TH_CNR = 0.2, TH_CM = 70%, and TH_por = 90% (definitions provided in **Appendix I, Section A**). OfUSA automatically generates a quality control report (**Table 1**), indicating the number of preserved and rejected trials. Using these criteria, 22% of visual trials and 18% of whisker trials were excluded. In the filtering step, the two principal components explaining the highest variance were removed to suppress residual global fluctuations. Each preprocessing step was conducted independently for data corresponding to the two stimuli.

**Table 1.** Accepted trials and rejection rates of all data as determined by quality control.

| AnimalID | SessionID | TaskID | Total Number | Accepted Trial Number | Rejected Trial Number | Reject Rate |
| --- | --- | --- | --- | --- | --- | --- |
| M221108 | 6IA4MV | V | 30 | 19 | 11 | 36.70% |
|  |  | W | 30 | 22 | 8 | 26.70% |
|  | 6OGYGK | V | 20 | 15 | 5 | 25.00% |
|  |  | W | 20 | 15 | 5 | 25.00% |
|  | 6R1GXD | V | 15 | 11 | 4 | 26.70% |
|  |  | W | 25 | 20 | 5 | 20.00% |
|  | 6RJFFT | V | 15 | 14 | 1 | 6.70% |
|  |  | W | 25 | 22 | 3 | 12.00% |
|  | 6SMC7V | V | 20 | 16 | 4 | 20.00% |
|  |  | W | 20 | 17 | 3 | 15.00% |
|  | 6UMOP5 | V | 20 | 19 | 1 | 5.00% |
|  |  | W | 20 | 20 | 0 | 0.00% |
|  | 6W74FL | V | 20 | 19 | 1 | 5.00% |
|  |  | W | 20 | 19 | 1 | 5.00% |
|  | 71U30T | V | 20 | 12 | 8 | 40.00% |
|  |  | W | 20 | 12 | 8 | 40.00% |
| Total |  | V | 160 | 125 | 35 | 21.88% |
|  |  | W | 180 | 147 | 33 | 18.33% |

Abbreviation: [V] denotes the visual stimulation, [W] denotes the whisker stimulation

The temporal information analysis component applies the “Segmentation ABA” process to perform region-by-time analyses. Using the regressors and ROI definitions specified during initiation, trial-specific time courses were extracted for each ROI (illustrated in **Figure 3b**). These trial-level time courses were then averaged across all trials within a session to generate a single representative time course per session (**Figure S1d**).

The spatial information analysis component employs a general linear model (GLM) at the voxel level to compute beta-value activation maps for each trial (illustrated in **Figure 3a**). Individual beta maps were then averaged across trials to yield a session-level activation map. Finally, this average beta map was transformed from the subject’s native space to the Allen Brain Atlas space using the pre-estimated transformation matrix.

#### 1.2.3 Group Level Analysis

In the group-level stage, random effects were estimated for both ROI time courses and activation maps by performing one-sample t-tests across sessions. The group-level analysis pipeline, illustrated in **Figure S1b**, is organized into three main components: initiation, temporal information analysis, and spatial information analysis.

The initiation component begins with the “Select Data” process, where the session-averaged time courses and activation maps obtained during the individual-level stage are selected for group analysis (via an interface similar to **Figure S1d)**. In the subsequent “Set Parameters” process, the relevant ROIs from the Allen Brain Atlas are specified, and labels are assigned to each experimental condition (e.g., “visual,” “whisker”).

For the temporal information analysis, the “Group T-test” process evaluates the significance of hemodynamic responses at each time point within every ROI. This is achieved by computing t-statistics across all sessions, thereby identifying time windows with consistent activation patterns. The outcomes are visualized using the “Barcode View” and “Time Course View,” which highlight temporal dynamics at the ROI level. **Figures 4c–d and S2a–d** illustrate representative region-by-time plots for visual cortical areas during stimulation tasks.

For the spatial information analysis, a t-test is applied to the beta maps across sessions to generate a group-level activation map, identifying brain regions showing statistically significant responses to the stimulus across the cohort. These maps are visualized using the “Mosaic View,” which produces multislice representations of the activation patterns (**Figures 4e and S2e–f**), and the “Regions Report,” which compiles a comprehensive summary of significantly activated brain regions (**Table 2**).

**Table 2.** Activated area of sensory stimulation in the rodent study.

| Stimulation | ROI_name | volume<br>(mm<sup>3</sup>) | Portion<br>(%) | Peak Value<br>T-Value | Peak coordinate(mm) |  |  |
| --- | --- | --- | --- | --- | --- | --- | --- |
|  |  |  |  |  | LR | AP | DV |
| Visual | SCi-L | 0.32 | 15.70 | 19.16 | -1.00 | 1.85 | -1.50 |
|  | SCs-L | 0.46 | 42.90 | 18.58 | -1.15 | 2.65 | -2.50 |
|  | VISI-L | 0.04 | 6.60 | 12.91 | -3.60 | 2.65 | -2.50 |
|  | VISp-L | 0.31 | 8.60 | 23.35 | -2.00 | 2.75 | -2.65 |
|  | VISpm-L | 0.04 | 7.10 | 12.39 | -1.40 | 1.85 | -2.85 |
|  | dLG-L | 0.04 | 10.70 | 12.42 | -2.35 | 1.15 | -0.95 |
|  | RSP-L | 0.37 | 7.10 | 27.42 | -1.45 | 2.95 | -2.70 |
| Whisker | AUDd-L | 0.14 | 22.60 | 13.66 | -4.40 | 0.45 | -1.10 |
|  | PO-L | 0.09 | 14.60 | 20.82 | -1.65 | 0.90 | -0.45 |
|  | SSp-bf-L | 0.95 | 30.10 | 29.74 | -3.40 | 0.25 | -1.45 |
|  | SSs-L | 0.48 | 10.50 | 18.40 | -3.70 | 0.80 | -2.00 |
|  | VP-L | 0.20 | 13.90 | 19.20 | -1.90 | 0.80 | -0.20 |
|  | SCi-L | 0.22 | 10.60 | 21.57 | -1.50 | 2.90 | -1.90 |
|  | SCs-L | 0.17 | 15.50 | 14.29 | -1.20 | 2.75 | -2.30 |
|  | VISrl-L | 0.04 | 8.10 | 14.00 | -2.60 | 0.65 | -2.85 |
|  | dLG-L | 0.05 | 12.70 | 10.26 | -2.15 | 1.10 | -0.60 |

### 1.3 Results

The group-level activation maps, together with the corresponding statistical reports, identified distinct brain regions showing significant responses to each sensory stimulus (one-sample t-test; T > 5.4, p < 0.001 uncorrected, df = 7, two-tailed). These results are presented in **Figures 4e** and **S2e–f**, with detailed information provided in **Table 2**.

Visual stimulation produced strong contralateral activation in regions well established for visual processing and sensory integration. Significant responses were observed in the Superior Colliculus (SCs, SCi), the primary and secondary visual cortices (VISp, VISl, VISpm), the Dorsal Lateral Geniculate nucleus (dLG), and the Retrosplenial cortex (RSP) (Hooks & Chen, 2020; Powell et al., 2020).

Whisker stimulation also evoked a robust contralateral response. Significant activation was detected along the canonical somatosensory pathway, including the primary and secondary somatosensory cortices (SSp-bf, SSs), the Ventral Posterior complex of the thalamus (VP), and the Posterior complex of the thalamus (PO). Beyond this core network, additional activation was observed in the Superior Colliculus (SCs, SCi), the Dorsal Lateral Geniculate nucleus (dLG), the rostrolateral visual area (VISrl), and the dorsal auditory cortex (AUDd). The recruitment of the primary somatosensory network aligns with the tactile nature of the whisker stimulus (Staiger & Petersen, 2021), while the involvement of subcortical structures such as the Superior Colliculus is consistent with evidence of their role in integrating non-visual sensory inputs (Benavidez et al., 2021). The additional activation observed in VISrl, AUDd, and dLG is less expected and may reflect experimental confounds, such as auditory noise generated by the stimulation device or visual detection of the whisker stimulator’s movement by the animal.

### 1.4 Summary

In summary, this demonstration presents a two-stage workflow for analyzing brain activity in response to visual and whisker stimulation, using a rodent atlas for spatial registration. First, data from each session is preprocessed at an individual level to extract hemodynamic response time courses and activation maps. Second, these metrics are consolidated for a group-level statistical analysis using t-tests. This method successfully revealed clear hemodynamic responses and activation maps in the relevant brain areas— specifically, visual stimuli activated the SCs and VISp, while whisker stimulation activated the SSp-bf and SSs.

## Application Example 2: Visual responses in an awake non-human primate

### 2.1 Experiment Setup

This study was conducted on a single male rhesus macaque trained to perform a passive fixation task for liquid reward. The animal was head-fixed and positioned in a sphinx posture, with its hands placed inside a box to reduce motion. During the task, the monkey fixated on a central point displayed on a high-resolution screen. While fixation was maintained, a large, colorful checkerboard stimulus was presented for 8 seconds at a flickering frequency of 10 Hz. Data were acquired in runs of approximately 18 minutes, each consisting of about 18 trials. **Figures 5a–b** illustrate the experimental setup and stimulus. Additional details are provided in **Appendix II, Section B**.

**Figure 5.**
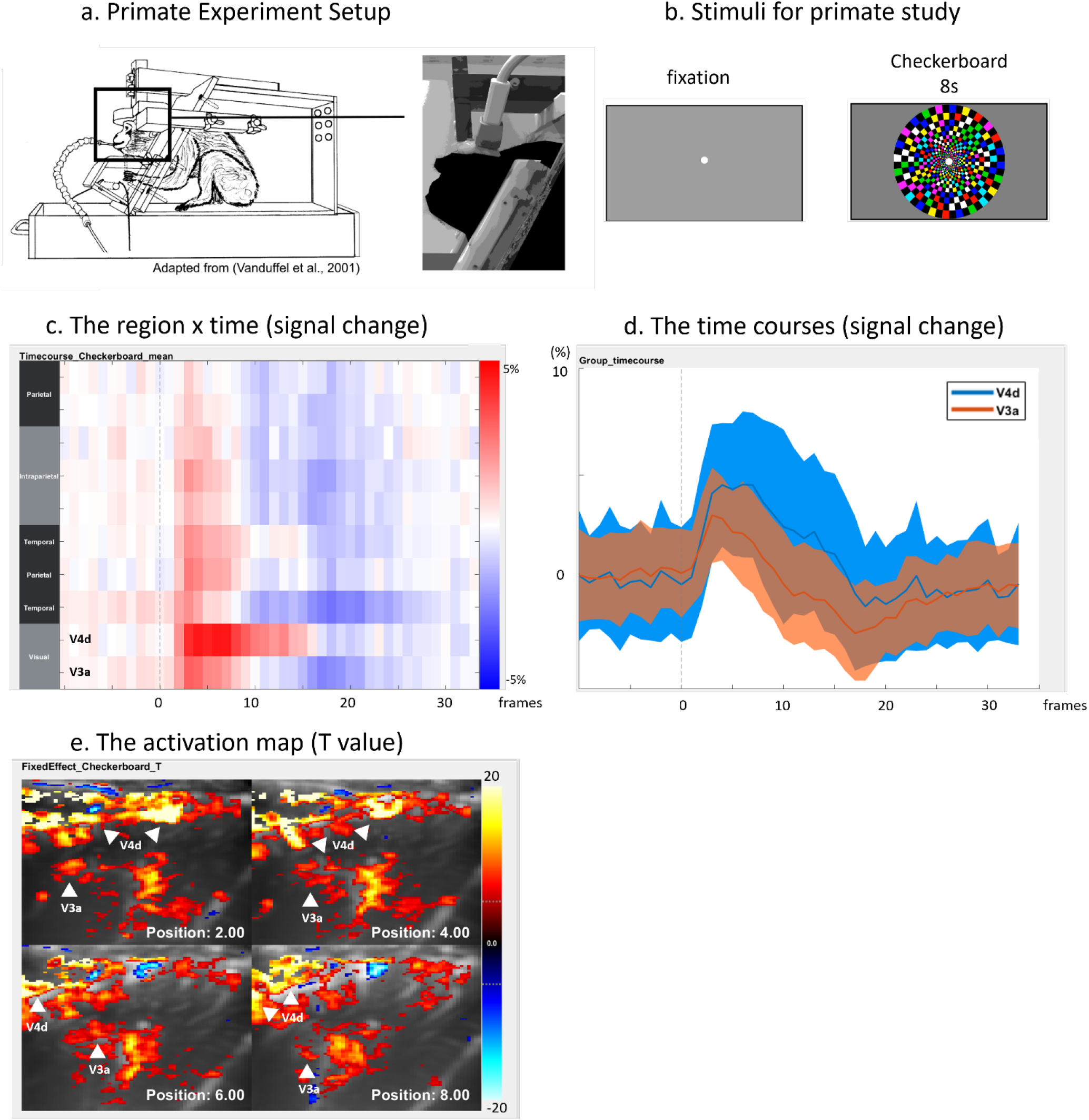
Experimental setups and results for the primate study. Panel (a) shows the setup for the visual experiment, in which the subject was surgically implanted with a headpost and a recording chamber to enable stable head fixation while seated in the sphinx position. The fUS transducer was mounted on the recording chamber for data acquisition. Panel (b) illustrates the visual task, consisting of an 8-second presentation of a colorful checkerboard on a 32-inch screen positioned 40 cm away, with the subject receiving rewards on a fixed schedule for maintaining fixation on a central point. Panel (c) presents region-by-time plots of time courses across multiple brain regions. Panel (d) displays representative time courses for visual regions (V4d, orange; V3a, blue). Panel (e) shows the group-level activation maps. Reported activations are statistically significant according to a fixed-effects analysis (T > 5.27, p < 0.05 FWE-corrected, df = 4850, two-tailed).

### 2.2 Analysis Workflow

Because no standardized fUS template exists for non-human primates, the analysis workflow was adapted to incorporate a custom study-specific template. This template was constructed by co-registering all sessions into a common anatomical space, enabling automatic alignment of individual datasets. The subsequent analysis followed the two-stage workflow described earlier. Given the limited number of sessions available, a fixed-effects model was employed at the group level, allowing the estimation of the average stimulus effect within the measured sample, rather than a generalizable random effect across sessions.

Trial structures for each run were specified in behavioral files formatted in JSON. These files contained detailed parameters for each stimulus event, including name, onset frame, duration, and expected time course. **Table S3** provides a complete description of these parameters.

#### 2.2.1 Study-Specific Template and ROI Definition

In the absence of a standard anatomical template for cross-session alignment, two strategies were evaluated. The first approach involved selecting a single session’s anatomical image as a reference. Although simple, this method risks introducing bias if the chosen session is not representative, potentially compromising the alignment of the remaining data. To overcome this limitation, a second and more robust strategy was adopted: generating a study-specific template by integrating anatomical images from all sessions. While more computationally demanding, this method reduces bias and improves cross-session registration accuracy.

The study-specific template was created using the “Data-driven Template” process, illustrated in **Figure S3a** and detailed in **Appendix I, Section B**. An initial template was generated by averaging anatomical images from all sessions. Each session’s image (**Figure S3b**) was then aligned to this template using the MATLAB imregtform function. The aligned images were averaged to form an updated template, and this cycle of registration and averaging was repeated ten times to obtain a refined consensus template (**Figure S3c**). Finally, the resulting template was manually registered to the CHARM Macaque Brain Atlas (Jung et al., 2021), enabling integration into a standard coordinate framework and facilitating the definition of regions of interest (ROIs) (**Figure S3d**).

#### 2.2.2 Individual-Level Analysis

The individual-level analysis pipeline for the primate dataset is illustrated in **Figure S3e**. Similar to the rodent workflow, it comprises four main components: initiation, preprocessing, temporal analysis, and spatial analysis.

The initiation component included two processes: “Loop Pipeline” and “Select Data.” In the “Loop Pipeline” process, the batch processing units (e.g., subject, session, stimulus) and their corresponding behavioral regressors were defined. In the “Select Data” process, the study-specific template (**Figure S3c**) and its associated regions of interest (ROIs) (**Figure S3d**) were specified for subsequent analyses.

The preprocessing component consisted of three steps: “Quality Control,” “Filtering,” and “Realignment.” Quality control was conducted using the same thresholds as in the rodent study to detect and exclude trials with burst errors. All datasets passed this stage, likely reflecting the effectiveness of pre-experimental training in reducing head motion. Filtering was then applied by removing the two principal components with the highest variance to suppress residual global fluctuations. Finally, the “Realignment” process automatically registered the functional data to the study-specific template. This step corrects for minor inter-session misalignments (up to ∼10% of the field of view) but is not designed to address severe distortions.

For temporal and spatial analyses, ROI time courses were extracted using the “Extract Timecourse” process and averaged across trials within each session using the “Average Across Data” process, yielding one representative time course per session. In parallel, a General Linear Model (GLM) was applied to each run through the “GLM Analysis” process, producing beta maps and corresponding variance maps. Because the GLM incorporated all trials within a session, each beta map represents the average stimulus-evoked response for that session and was subsequently used in the group-level analysis.

#### 2.2.3 Group-Level Analysis

At the group level, a fixed-effects analysis was performed to calculate the average ROI time courses and activation maps across all sessions. The group-level pipeline, shown in **Figure S3f**, included three primary components: initiation, temporal analysis, and spatial analysis.

The initiation stage consisted of three “Select Data” processes and one “Set Parameters” process. The first three processes specified the session-level ROI time courses, the corresponding activation maps, and the study-specific template with its ROIs. In the “Set Parameters” process, the experimental condition was labeled as “checkerboard.”

For temporal analysis, the “Average Across Data” process was used to compute the mean hemodynamic response across all sessions for each ROI and time point. These responses were visualized with the “Barcode View” and “Time Course View,” enabling a clear representation of stimulus-evoked activity. **Figures 5c–d** show the resulting region-by-time plots summarizing the average response to the checkerboard stimulus.

For spatial analysis, the “Fixed Effect Analysis” process combined the beta maps from all sessions to generate a group-level activation map. This map identifies brain regions consistently activated by the stimulus across the dataset. The final results were visualized using the “Mosaic View” to produce multislice representations of activation (**Figure 5e**) and the “Regions Report” process to compile a summary table of significantly activated regions (**Table 3**).

**Table 3.**
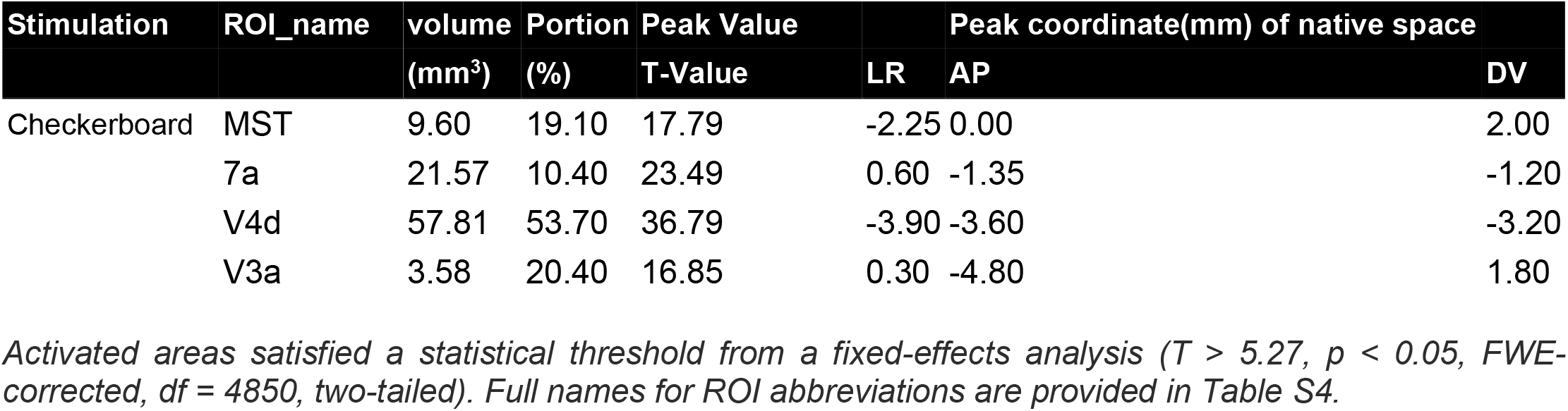
Activated area of checkerboard stimulation in the monkey study.

### 2.3 Results

The group-level activation maps and ROI time courses revealed distinct regions of significant activation in response to the checkerboard stimulus. Results are shown in **Figures 5c–e** and **Table 3**. Statistical thresholds were set at p < 0.05, Family-Wise Error (FWE) corrected (fixed-effects; T > 5.27, df = 4850, two-tailed).

Quantitative analysis identified the strongest responses in visual areas V4d and V3a, which exhibited signal changes of approximately 3–5% and maximum T-values of 36.78 and 16.85, respectively. Additional significant activation was observed in areas MST and 7a, showing signal changes of around 2% with maximum T-values of 17.79 and 23.49, respectively.

These findings are consistent with previous reports describing the involvement of both ventral and dorsal visual streams in processing structured visual stimuli, in particular checkerboard-like patterns that drive responses in areas V4d, V3a, 7a, and MST (Arcaro & Livingstone, 2017; D. C. Van Essen et al., 1992; David C. Van Essen & Maunsell, 1983). Robust activation in V4d, a core region of the ventral stream, aligns with its established role in processing contrast and edges. Engagement of the dorsal stream was reflected in V3a, which contributes to processing global visual forms, and in area 7a, which integrates visual information for visuomotor functions. By contrast, the weaker activation in MST is consistent with its specialization in optic flow processing, which was not a primary feature of the stimulus.

Overall, the results reveal a dominant involvement of the ventral visual stream, associated with object recognition, and a more transient contribution of the dorsal stream, consistent with its role in dynamic visual integration.

### 2.4 Summary

This demonstration presents a full workflow for analyzing stimulus-evoked activity in the primate visual cortex. The approach begins with the construction of a study-specific template to achieve robust alignment across sessions, followed by a two-stage statistical procedure. At the first level, individual analyses extract hemodynamic time courses and activation maps from preprocessed data. At the second level, a group fixed-effects analysis integrates these outputs to reveal consistent activation patterns. The results clearly demonstrate stimulus-driven activation in key regions of the ventral (V4d) and dorsal (V3a) visual streams.

## Rationale and Justification of the Proposed Workflow

### 3.1 Advantages

This paper introduces OfUSA, an open-source and open-access software package for the analysis of functional ultrasound (fUS) data. OfUSA addresses critical methodological needs in the fUS research community and provides several distinctive advantages.

First, OfUSA is designed for accessibility. It incorporates a comprehensive graphical user interface (GUI) that makes the software usable by researchers with or without programming experience. As illustrated in Figures 2 and S1c–e, the GUI streamlines essential tasks such as data management, image registration, and the creation of experimental regressors. The visualization interface further enables users to interactively adjust parameters, generate figures, and apply statistical thresholds with ease.

Second, OfUSA establishes a standardized yet flexible workflow. The pipeline covers the entire analytical process, from preprocessing to statistical analysis and visualization, ensuring consistency and reproducibility. Built-in statistical tools support both spatial activation mapping and temporal hemodynamic response analysis at individual and group levels. While this framework enforces methodological rigor, it also allows parameter adjustments (e.g., thresholds, preprocessing settings) to accommodate diverse experimental paradigms. Its batch-processing capacity further enables efficient analysis of large-scale datasets across sessions and animals.

Third, OfUSA prioritizes compatibility and extensibility. Data are stored in widely adopted NIFTI and JSON formats, ensuring interoperability with established third-party tools. As an open-source project, OfUSA also invites contributions from the scientific community, encouraging integration of new methods and continuous evolution of the software in parallel with advances in the field.

Fourth, OfUSA provides advanced visualization capabilities. A set of integrated toolboxes (Section 3.5) enables researchers to explore and communicate findings using multiple complementary perspectives on brain activity, including spatial, temporal, and spatiotemporal representations.

Together, these advantages establish OfUSA as a complete, end-to-end solution for modern fUS research. The following sections further justify the methodological choices of the workflow, including preprocessing, modeling of the hemodynamic response, fixed-versus random-effects analyses, visualization strategies, and current limitations.

### 3.2 Preprocessing

Preprocessing is a critical step in fUS analysis, aimed at reducing noise and minimizing confounding influences. Although fUS measures cerebral blood volume (CBV), insights can be drawn from fMRI studies of the BOLD signal, where motion (48%) and physiological fluctuations (31%) account for a large portion of variance (Liu et al., 2017).

A unique challenge in fUS is burst noise, typically arising from motion, which can exceed baseline signal amplitude by several orders of magnitude (Brunner et al., 2021). This can be mitigated behaviorally, through animal training to reduce head movement (as shown in our primate study), and computationally, through OfUSA’s preprocessing steps. Quality control and filtering are central to this process: quality control identifies and rejects corrupted trials, while filtering suppresses residual global fluctuations.

To evaluate preprocessing strategies, four conditions were compared: standard (quality control and filtering), no preprocessing, quality control only, and filtering only. Analyses of activation maps and time courses (**Figures S4–S5**) revealed that the standard pipeline reduced spurious activations and minimized cross-trial variance, though with slightly lower raw signal amplitude. Filtering without prior quality control induced negative artifacts, a well-documented risk when global components are removed in the presence of strong burst noise (Aguirre et al., 1998). These results confirm that filtering is most effective when preceded by quality control.

Performance was further quantified using SNR, CNR, and d-prime (d ′), revealing species-dependent effects. The standard pipeline produced significantly higher SNR in rodents, while QC only provided meaningful improvements in primates but not in rodents. Filtering alone degraded metrics in rodents while partially improving CNR and d ′ in primates, likely reflecting differences in burst noise prevalence.

The choice of preprocessing strategy significantly impacts fUS data quality, with the most robust approach being a combination of quality control (QC) and filtering. Since optimal parameters vary by experiment, the following guidelines should serve as a starting point. For the QC step, recommended ranges are a burst error threshold between 200-400%, an SNR threshold between 5-10, and a CNR threshold between 0.5-2. Applying stricter criteria—such as a burst error threshold below 200% or using higher SNR/CNR thresholds— selects for better quality data. This enhances statistical power, meaning fewer trials are needed to detect the same signal change. However, the trade-off is that stricter criteria may also lead to a high rejection rate of trials. In contrast, a relatively relaxed common noise threshold (0.6 to 0.8) is sufficient, as the subsequent filtering process will effectively remove residual common mode effects.

During the PCA-based filtering process, we recommend removing up to three principal components. The effectiveness of this choice can be easily diagnosed. If the resulting activation map shows widespread negative activations, it is likely that too many components have been removed. Conversely, if a region-by-time plot reveals highly synchronized patterns across the brain, often appearing as vertical stripes, it suggests too few components were removed and residual common noise persists. These parameters should be adjusted based on such diagnostic checks to best suit the specific characteristics of your dataset.

### 3.3 Hemodynamic Response Function

OfUSA provides tools to estimate hemodynamic response functions (HRFs) across brain regions, leveraging fUS’s high spatial and temporal resolution. Results (**Figures 4d, 5d, S4b, S5b**) demonstrate that HRFs vary substantially between regions, consistent with prior work (Brunner et al., 2020; Lambert, Niknejad, et al., 2025; É. Macé et al., 2018; Sans-Dublanc et al., 2021).

This heterogeneity challenges conventional fMRI approaches that assume a uniform canonical HRF. Such assumptions risk underestimating activation strength, inflating error rates, and biasing connectivity estimates (Handwerker et al., 2012, 2004; Rangaprakash et al., 2018). OfUSA avoids this by employing paradigm-based regressors in GLM analyses without convolving with a canonical HRF, yielding direct and unbiased activation estimates. Future extensions could integrate empirical HRF modeling to further refine region-specific neurovascular dynamics.

### 3.4 Fixed versus Random Effects

Both fixed- and random-effects models are implemented in OfUSA. Random-effects models generalize findings across populations by accounting for variability across subjects or sessions, as demonstrated in our rodent dataset. Fixed-effects models, in contrast, assess the average effect within the measured sample only, as shown in the primate dataset.

While random-effects models offer broader generalizability, their adoption in fUS research is limited by the logistical challenges of large sample sizes, given the surgical and training demands of animal preparation. For smaller cohorts, fixed-effects analyses remain appropriate. As fUS adoption grows and larger datasets become available, increased use of random-effects models is anticipated.

### 3.5 Visualization

OfUSA integrates four visualization tools: multislice (mosaic) view, 3D rendering, region-time view, and time-course view. These tools allow flexible exploration of spatial, temporal, and spatiotemporal data.

The multislice view generates brain-wide activation maps overlaid on reference templates or native-space images. Reports include peak strength, coordinates, and activation volume (**Tables 2–3**). The 3D rendering tool, currently implemented for the mouse brain, projects activation strength onto brain surfaces, with optional ROI overlays for contextual interpretation (**Video S1-S2**).

The region-time view presents barcode-like spatiotemporal plots of regional responses, while the time-course view enables interactive exploration of response dynamics in single or multiple ROIs. Both tools support customizable ROI selection and ordering.

These visualization options provide complementary insights but require careful parameter choices, particularly for thresholds and color scales. To ensure transparency and reproducibility, all visualization parameters should be reported alongside figures.

### 3.6 Limitations

The current version of OfUSA has limitations that highlight directions for future improvement.

Firstly, it currently relies on manual registration and lacks a tool for correcting brain distortion, as robust automated registration algorithms are not yet widely available. However, OfUSA’s modular architecture is designed to seamlessly integrate such tools once they are developed. Secondly, the quality control process requires manual threshold selection because optimal parameters vary with the experimental setup and paradigm. This limitation is expected to be resolved as the growing adoption of fUS technology provides the large datasets needed to build more sophisticated and adaptive methods. Finally, limitations inherited from its MATLAB environment, such as license costs and potential for slow performance, are also addressed. The cost barrier is lowered by ensuring compatibility with the free MATLAB Runtime environment, while performance is continuously improved through code optimization and the option to use the Parallel Computing Toolbox.

## Conclusions

OfUSA provides a unified, open-source platform for preprocessing, analyzing, and visualizing fUS data. Its GUI-based interface lowers the barrier for researchers without programming expertise while preserving methodological rigor and flexibility. Demonstrations on rodent and primate datasets highlight its strengths in preprocessing, spatial and temporal analysis, and visualization.

By addressing critical analytical bottlenecks, OfUSA supports broader adoption of fUS in neuroscience. It reduces the reliance on specialized coding expertise, facilitates reproducible workflows, and enables researchers to explore whole-brain activity in awake animals with greater efficiency. As part of the OpenfUS initiative, OfUSA represents a major step toward accelerating collaborative, large-scale, and translational fUS research.

## Data availability

Dataset used in this work are available on a Zenodo repository at: https://doi.org/10.5281/zenodo.17130250

## Code availability

OfUSA software can be found and downloaded from the GitHub repository: https://github.com/YunAnGitHub/OpenfUS_Analyzer_OfUSA.

For detailed instructions and guidance, please refer to the user manual available at our Help Center: https://openfus.notion.site/OpenfUS-Analyzer-Help-Center-fec9cf928efc4e0eb257e0f24c29b992

## License

The software is freely available for educational and research use under the CC BY-NC-SA 4.0 license. For details, please see: https://creativecommons.org/licenses/by-nc-sa/4.0/

## Acknowledgements

This work is supported by grants from Fonds Wetenschappelijk Onderzoek (G079623N, G091719N, 1197818N, G0A4F24N, G055124N), ERANET, EU Horizon 2020 (Grant number 964215, UnscrAMBLY), HORIZON-MSCA-2022-DN-01 (Project 101119916-SOPRANI).

We are grateful to Dr. Mathilda Froesel for her support with the monkey experiments.

## Contributions

Y.A. Huang and A. Urban conceived the work; M. Grillet acquired the rodent data; N.E. Fitzgerald and M. Froesel designed the experiment and acquired data of primate study; W. Vanduffel supervised the primate experiment; Y.A. Huang developed the software; Y.A. Huang, T. Lambert and D. Verbeyst designed the framework; Y.A. Huang and C. Brunner developed the standardized workflow; Y.A. Huang and N.E. Fitzgerald developed the analysis workflow for primate study; Y.A. Huang wrote the manuscript under the supervision of A. Urban; All authors discussed the results and contributed to the final manuscript.

## Declaration of generative AI and AI-assisted technologies in the writing process

During the preparation of this manuscript, the authors used generative AI tools (Gemini and ChatGPT) to assist with grammatical corrections and to improve readability and language clarity. All content generated with these tools was subsequently reviewed and edited by the authors, who take full responsibility for the final version of the published article.

## Appendix I

### A. Process of the standardized workflow

This standardized analyzing workflow that was adapted from our previously published procedures (Brunner et al., 2021; É. Macé et al., 2018). The workflow comprises two stages, the individual stage and group stage. In the former stage, we process all the data independently (trial-by-trial or session-by-session). Specifically, we preprocess the data and analyze the data for generating the activation map and extracting the ROIs’ hemodynamic time course. In the latter stage, we group all the individual results together for generating the group-level activation map and time courses of ROIs. We also visualize the final results in this stage. In the following section, we introduce the formula of all process we used in the standardized analyzing workflow.

#### a. Individual Stage

##### a.1. Preprocessing

###### a.1.1. Quality Control

Quality control consists of four criteria: burst error, temporal Signal-to-Noise Ratio (tSNR), Contrast-to-Noise Ratio, and common mode.

###### a.1.1.1. Burst Error

An artifact commonly observed in functional ultrasound imaging is the burst error. This artifact manifests as a sudden and large signal across the image, primarily caused by the movement of the imaged object. To measure the burst error, we use the ratio of the peak value to the average baseline signal in the scan period. Mathematically, this can be expressed as:

I(x,y,z,t) denotes the 4D images, where I:ℝ^*X×Y×Z×T*^, x ∈ *X*, y ∈ Y, z ∈ Z, t ∈ T. X, Y, Z, and T represent the range of images in the X, Y, Z directions, and the total length of timepoints, respectively.

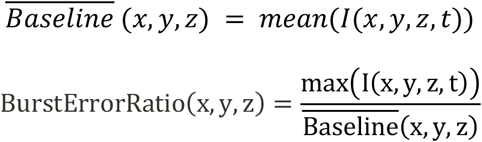

To control the burst error, we have introduced two thresholds: the burst error threshold (TH_bursterror_) and the noisy voxel threshold (TH_noisyvoxels_). The TH_bursterror_ is used to estimate the voxels affected by the burst error, while the TH_noisyvoxels_ is used to estimate the cover rate of burst error voxels in a trial. By applying a mask to remove the noisy voxels, we accept the trials only if the number of noisy voxels does not exceed the TH_noisyvoxels_. The mask and the accepted trials can be represented as follows:

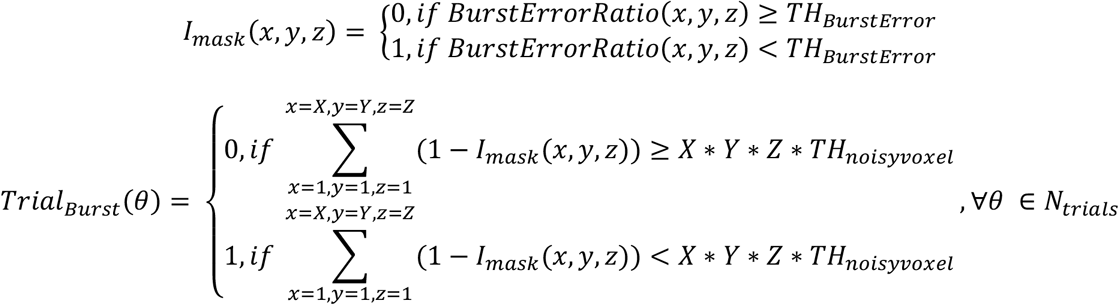

where *N*_*trials*_ denotes the total number of trials.

###### a.1.1.2. temporal Signal-to-Noise Ratio (tSNR)

The tSNR (temporal signal-to-noise ratio) is a useful indicator for estimating the amount of noise fluctuation relative to the signal. In general, a higher level of noise leads to a lower tSNR. To obtain a reliable signal, researchers often need to increase the number of trials to better control the noise level in the group stage. Therefore, tSNR helps researchers estimate the minimum number of trials required to ensure the reliability of the measured signal.

The global tSNR is calculated by dividing the standard deviation of baseline fluctuation (noise) by the average intensity of the signal (signal) for each voxel. Mathematically, it can be expressed as:

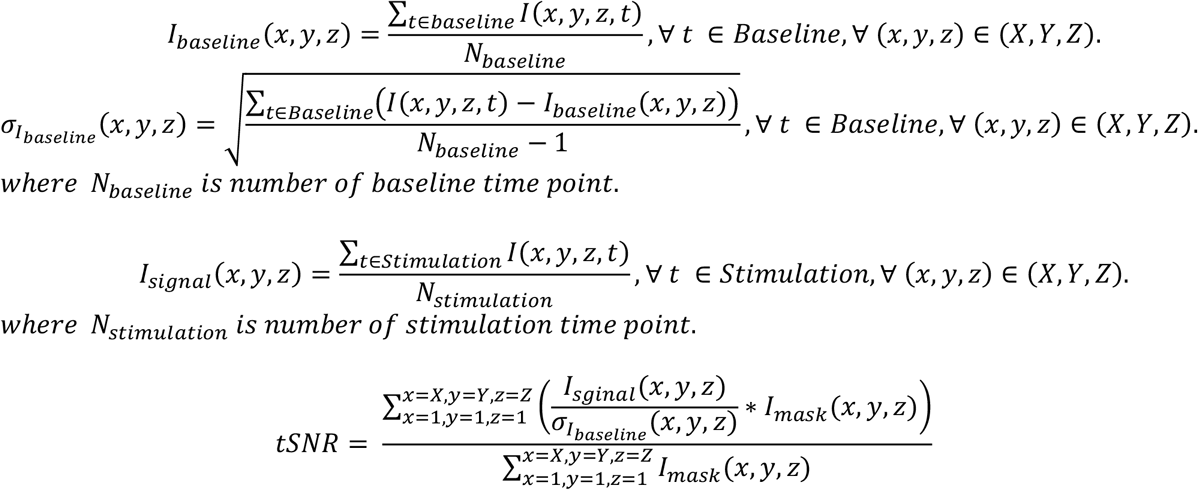

Notably, we computed the global tSNR without including the noisy voxels identified through the burst error criteria. This allowed us to obtain a more accurate estimation of the true tSNR, unaffected by burst errors.

To ensure that the data was not influenced by high levels of noise, we introduced a signal-to-noise ratio threshold (TH_SNR_). Trials were deemed valid if the tSNR value exceeded this threshold. This can be expressed as follows:

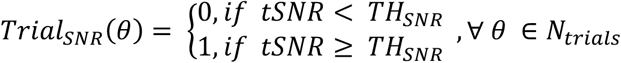

###### a.1.1.3. Contrast-to-Noise Ratio (CNR)

The CNR (Contrast-to-Noise Ratio) is an indicator that measures the ratio between the signal change associated with stimulation and the noise level. While the tSNR reflects the reliability of the signal, the CNR reflects the reliability of the measured signal change. Furthermore, the CNR is directly related to the probability of observing clear hemodynamic curves from the time course

The global CNR is estimated by considering the standard deviation of baseline fluctuation (noise), the average intensity of the stimulation signal (signal), and the average intensity of the baseline signal (baseline) for each voxel. Mathematically, it can be expressed as:

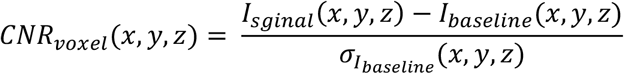

Since not all voxels are associated with the given stimuli, we consider the CNR from the top percentage (P_CNR_) of voxels, which can be described as the percentile of CNR.

Since not all voxels are associated with the given stimuli, we consider the CNR from the top *percentage* (P_CNR_) of voxels, which can be described as the *percentile* of CNR (P_CNR_).

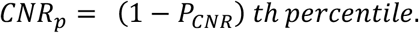

For example, P_CNR_ = 5% means that the *top 5%* of CNR voxels would be considered. CNRp denotes the *95*^*th*^ *percentile* of CNR. Furthermore, we can define the global CNR and the accepted trials with a CNR threshold (TH_CNR_) as follows:

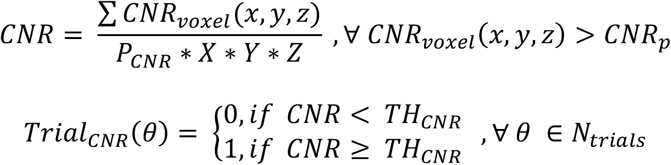

###### a.1.1.4. Common Mode

The common mode refers to the presence of correlated fluctuations throughout the brain, such as those caused by respiratory rate, heart rate, pupil size, and awareness. These fluctuations can also be caused by confounding factors such as movements or system noise from the imaging technique. While global signal filtering can be used to remove common fluctuations after quality control, this filtering is only effective for moderate levels of common mode fluctuation. Therefore, it is crucial for researchers to be aware of the level of common mode fluctuation in their data. If trials contain severe common mode fluctuation, they should be excluded from analysis because the global signal filtering doesn’t work well in this case.

To efficiently estimate common mode fluctuation, we propose a rapid and computationally efficient method. We divide the entire brain volume into smaller subblocks of equal size, designated as Nblock x Nblock x Nblock. For instance, Nblock = 10 represents a grid size of 10, resulting in a total of 1000 subblocks. Next, we calculate the pairwise correlation between these subblocks and estimate the common mode level using two thresholds: the common mode correlation threshold (TH_CM_) and the common mode portion threshold (TH_por_). The formula for this calculation is as follows:

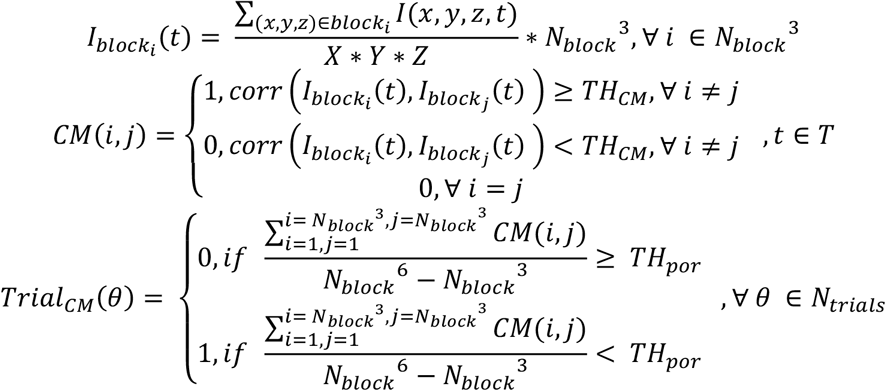

where *corr*(*f*(*t*), *g(t*)) refers to the Pearson’ s correlation coefficient of timecourse *f(t) and g*(*t*).

###### a.1.1.5. Integration of four criteria

Finally, a trial would be deemed acceptable if it fulfills all four criteria. Accepted trials would be listed as follows:

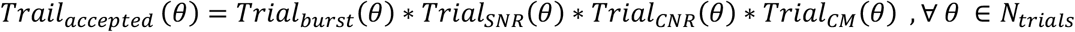

###### a.1.2. Filtering

To eliminate the common mode fluctuation described earlier, we implemented a global signal filtering method based on principal component analysis (PCA). Since the global fluctuations are significantly larger than the hemodynamic response, removing these global fluctuations allows us to uncover the underlying activation patterns associated with local neural activity. We constructed the common mode fluctuation from the first Nth components and subsequently subtracted this common mode fluctuation from the time course of each voxel.

To optimize computational efficiency, we first transform the 4D image I(x, y, z, t) into a 2D matrix I(m, t) using a rearrangement function 𝒯.

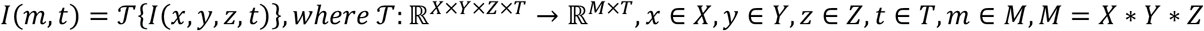

Utilizing the built-in principal component analysis (PCA) function *pca* in MATLAB, we can represent I(m, t) as the product of the score matrix (S) and coefficient matrix I.

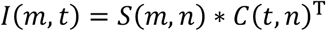

where *I*: ℝ^*M*×*T*^, *S*: ℝ^*M*×*N*^, *C*: ℝ^*T×N*^, *N* is the number of components, T denotes transpose of a matrix Denoting the signal reconstructed from the first Nth components as PCA_N_, we can formulate the filtered data by subtracting PCA_N_ from the original data:

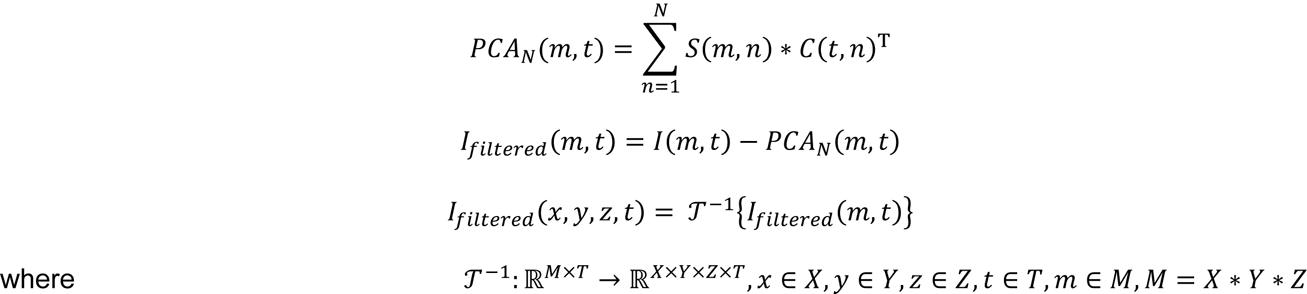

###### a.1.3. Registration

The final step in the preprocessing stage involves estimating the transformation matrix to align the brain volume from native space to the Allen Brain Atlas Common Coordinate Framework (ABA CCF). Researchers can utilize a graphical user interface (GUI) to estimate the nine parameters of the affine transformation matrix. These parameters encompass the shifts in left-right (LR), anterior-posterior (AP), and dorsal-ventral (DV) directions, rotations along the LR, AP, and DV axes, and scaling along the LR, AP, and DV directions. This registration interface, initially introduced in our previous publications (Brunner et al., 2021), has been seamlessly integrated into the OfUSA framework.

The registration process 𝒯_*ABA*_ and the resulting affine transformation matrix *T*_*ABA*_ can be represented by the following formula:

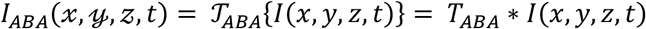

where 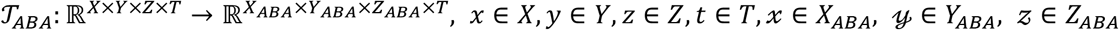. *X, Y, Z* denotes the native space and *X*_*ABA*_, *Y*_*ABA*_, *Z*_*ABA*_ denotes Allen Brain Atlas space.

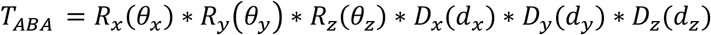

where each of rotation and transition matrix could be written as:

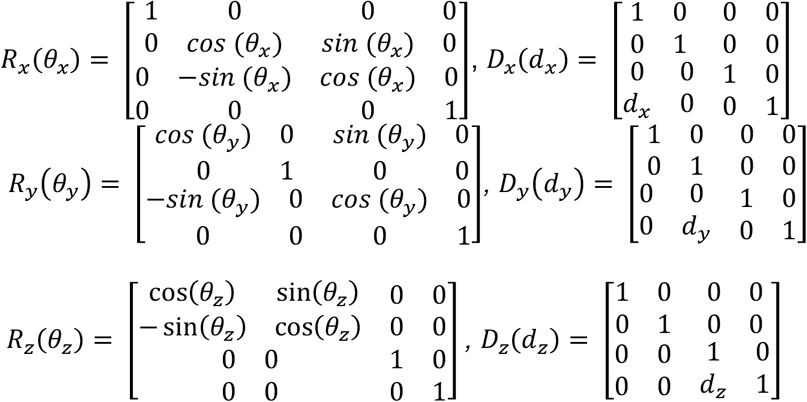

The matrix *R*_*x*_(*θ*_*x*_), *R*_*y*_(*θ*_*y*_), *R*_*z*_(*θ*_*z*_), *D*_*x*_(*d*_*x*_), *D*_*y*_(*d*_*y*_), and *D*_*z*_(*d*_*z*_) can be obtained by employing the provided registration interface to manually determine the rotation angle and transition level.

##### a.2. Individual-level analysis

###### a.2.1. Activation map analysis

To determine the activation map, we employed the general linear model (GLM) to quantify the signal change elicited by the stimulus in each voxel. By evaluating the signal change across all voxels in the brain, we generated an activation map representing the spatial distribution of stimulus response.

The time course of a voxel is denoted by *I*_*x,y,z*_(*t*) and can be represented as the sum of the stimulus-related activation (I_reg_), error term I, and a constant baseline bias I (Friston et al., 1994):

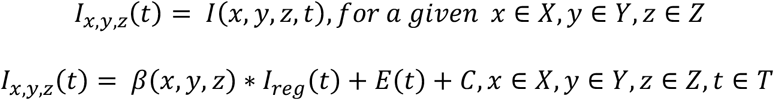

where *β* denotes the signal of each voxel and *I*_*reg*_ denotes the ideal activation pattern.

The *β*(*x, y, z*) represents the signal change of the response in a voxel to a specific stimulus condition. It is calculated by solving the linear regression equation. Define the Q matrix by combining the I_reg_ and the constant term C.

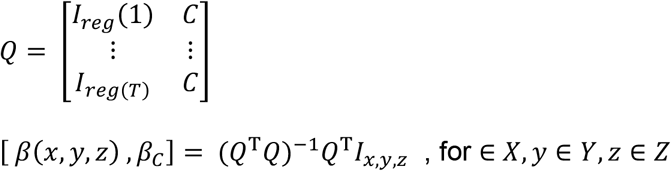

where *β*_*C*_ denotes the baseline bias, T denotes transpose of a matrix, and *Q*: ℝ^*T*×2^, *I*_*x,y,z*_: ℝ^*T*×1^

The variance matrix, denoted as *Var*(*x, y, z*), represents the variance in the GLM analysis for a given voxel. It’s calculated using mean square error (MSE) and degree of freedom (DOF) from the analysis.

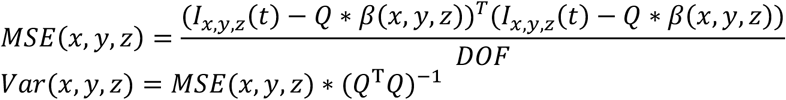

It is important to note that the GLM can accommodate multiple regressors, including multiple stimuli or other confounding factors such as animal movements. However, due to the simplicity of the paradigm in this study, we did not utilize these additional regressors. This functionality could be implemented in the future. Secondly, unlike fMRI studies that often employ an ideal paradigm convolved with an impulse response function, also known as the canonical hemodynamic response function, derived from the gamma function (Penny et al., 2007), we did not implement any impulse response function in this study. This is because the shape of the hemodynamic response exhibits significant variation across regions and stimuli. While the impulse response function is an important concept, measuring the empirical impulse response function for each region and stimulus is not the primary objective of this study. Moreover, incorporating the empirical impulse response function would have only a limited impact on the simple paradigm we demonstrate in this work.

Once we obtained the activation map *β*(*x, y, z*), we mapped it to the Allen Brain Atlas (ABA) Space using the transformation matrix obtained in the preprocessing step.

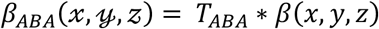

where *x* ∈ *X, y* ∈ *Y, z* ∈ *Z, t* ∈ *T, x* ∈ *X*_*ABA*_, *𝓎* ∈ *Y*_*ABA*_, *𝓏* ∈ Z_*ABA*_. This step was performed using the built-in MATLAB function imwarp.

###### a.2.2. Region-time analysis

To efficiently compute the time course of different regions, we developed an algorithm that minimizes computational burden. Instead of the computationally demanding approach of wrapping the entire 4D matrix I(x, y, z, t) to the Allen Brain Atlas (ABA) space and averaging the signal of voxels within regions, our algorithm directly calculates the coordinates of the regions in the native space. This is achieved using the inverse transform function, *transformPointsInverse*, built-in in MATLAB, along with linear interpolation to align the inverse transform function with the native space. By directly computing the time course of each region in its native space, we significantly reduce computational demands while maintaining accurate time course measurements.

We denote the 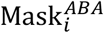 as the voxels of i^th^ regions of Allen Brain Atlas and denote 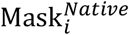 as the voxels of i^th^ regions of Allen Brain Atlas in the native space. The measurement of time course of i^th^ region (I_i_(*t*)) can be written as:

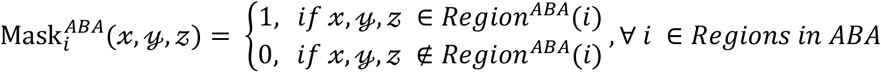

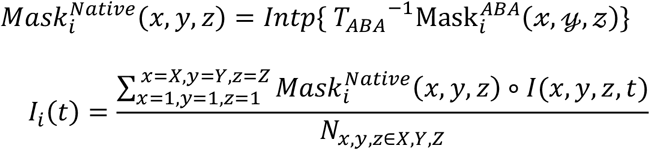

Where *𝓍* ∈ *X*_*ABA*_, *𝓎* ∈ *Y*_*ABA*_, *𝓏* ∈ Z_*ABA*_, *x* ∈ *X, y* ∈ *Y, z* ∈ *Z, t ∈ T, T*_*ABA*_^−1^ denotes inverse transform function, Int*p* denotes the trilinear interpolation, *“* ∘” denotes Hadamard product (element-wise product), and *N*_*x,y,z*∈*X,Y,Z*_ denotes the number of voxels in i^th^ region of Allen Brain Atlas in the native space.

Since the coordinates obtained from the inverse transform are not always whole numbers, the interpolation function (Intp) is employed to map the intensity values from the non-integer coordinates to the nearest whole-number coordinates (original native coordinates). This interpolation process can be mathematically expressed as follows:

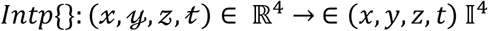

where ℝ^4^ *and* 𝕀^4^ represent the real numbers and integer of four-dimensional coordinates space, respectively.

#### b. Group Stage

##### b.1. Group-Level Analysis

After obtaining the activation maps or region-time analysis on the individual level. We can investigate the group effects across these measures. To this end, we introduce two methods: averaging and t-test. Both methods aid researchers in comprehending the common patterns within their data. Averaging summarizes a set of data, while the t-test provides statistical inference for the hypothesized effects. This allows researchers to study both of the fixed effect (averaged based on part of population) or random effect (statistical inference on the entire population) of the given stimuli.

###### b.1.1. Averaging for fixed effect

Averaging is a powerful tool for summarizing a set of data. We offer two types of averaging methods: arithmetic mean (default) and median mean. The arithmetic mean is the most commonly used method, but it is not resistant to extreme value. If the dataset contains extreme value, the median value, which represents the middle value of the dataset, provides a more reliable summary. Both arithmetic mean and median can be applied to activation maps or region-time data.

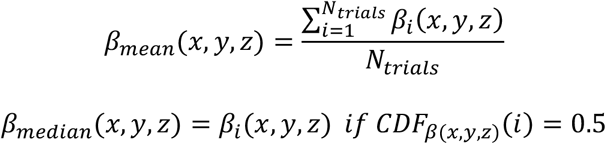

Where *β*_*i*_ denotes the activation map of i^th^ trials, and *x* ∈ *X, y* ∈ *Y, z* ∈ *Z. CDF*_*β*(*x,y,z*)_ represents the cumulative distribution function of activation maps at coordinate *x, y, z*.

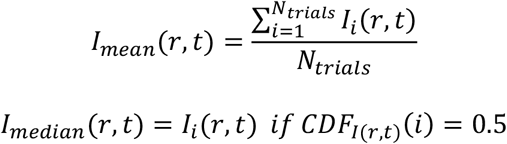

where *I*_*i*_(*r, t*) denotes the region-time matrix of i^th^ trials, and r ∈ *N*_*ABA*_, *t ∈ T, N*_*ABA*_ denotes the regions in Allen Brain Atlas space. *CDF*_*I*(*r,t*)_ represents the cumulative distribution function of region-time intensity of r^th^ region at timepoint t.

###### b.1.2. Joint Beta value and T value for fixed effect analysis

To calculate the fixed effect, we first combine the individual beta values and their variances. The overall T-value is then computed from these pooled estimates, as shown below:

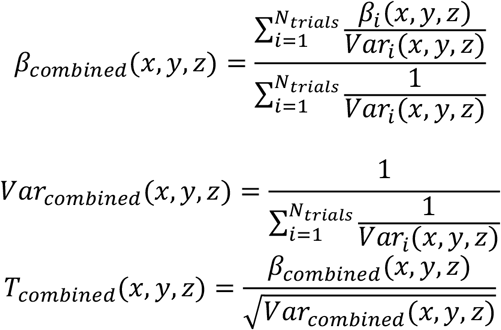

Where *β*_*i*_(*x, y, z*) denotes the activation map of i^th^ trials and *Var*_*i*_(*x, y, z*) denotes the variance map of i^th^ trials.

###### b.1.3. T-test for random effect

The t-test is a widely used statistical method for hypothesis testing (alternative hypothesis). For instance, it can be used to determine whether the activation in a particular region is significantly higher than the baseline. In this case, the null hypothesis assumes that the activation signal change is equal to the baseline. Alternatively, it can be employed to test the alternative hypothesis of a difference between two groups (alternative hypothesis) by rejecting the null hypothesis that the signal change of the two groups is equal. We have implemented one-sample t-tests, two-sample t-tests, and paired t-tests, allowing researchers to select the appropriate method based on their data and the research question they aim to address.

The t-test can be applied to both activation maps and region-time data. The activation map and region-time matrix of one-sample t-tests can be written as follows:

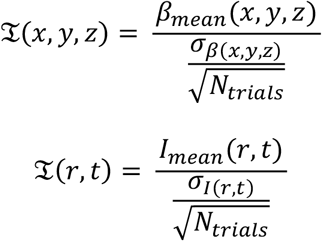

where *x* ∈ *X, y* ∈ *Y, z ∈ Z, t ∈ T*. 𝔗(*x, y, z*) denotes the t-value of activation strength of (x,y,z) coordinates, and 𝔗(*r, t*) denotes the t-value of activation strength of rth region at timepoint t. *σ*_*β*(*x,y,z*)_ and *σ*_*I*(*r,t*)_ denotes the standard deviation of the activation maps at coordinate (*x, y, z*) and intensity of r^th^ region at timepoint t, respectively, across trials. It can be written as:

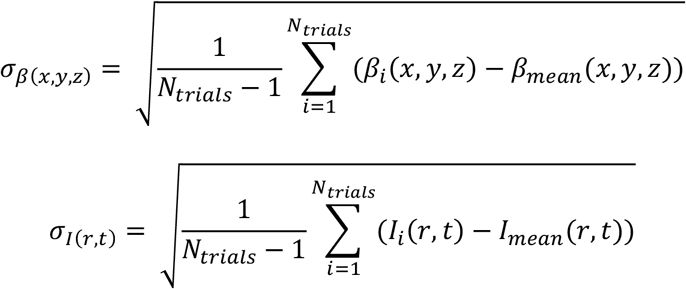

Similarly, the two-sample t-test can be written:

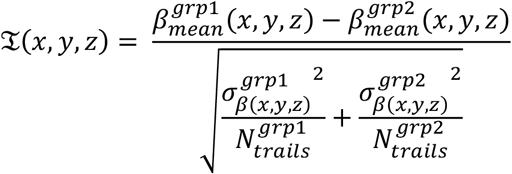

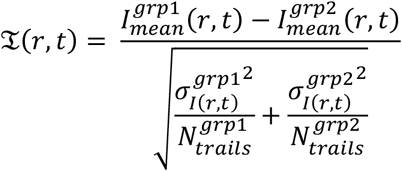

where *x* ∈ *X, y* ∈ *Y, z* ∈ *Z, t ∈ T*, 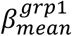 and 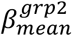 denotes the mean activation at coordinate (x,y,z) of group 1 and group 2, respectively. 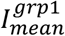 and 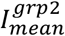 denotes the mean intensity of r^th^ reigons at timepoint t of group 1 and group 2, respectively. Similarly, the 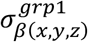 and 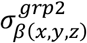 denotes the standard deviation of the activation at coordinate (x,y,z) of group 1 and group 2 across the trials, respectively. 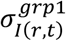 and 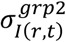 denotes the standard deviation of the intensity of r^th^ reigons at timepoint t across trials of group 1 and group 2, respectively. 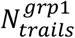 and 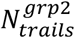 denotes the trials number of group 1 and group 2, respectively. The sample number in two groups is not necessarily equal.

The paired t-test is employed to determine whether the samples measured at different time points, denoted as group 1 and group 2, exhibit significant differences. The paired t-test can be directly applied to the activation map:

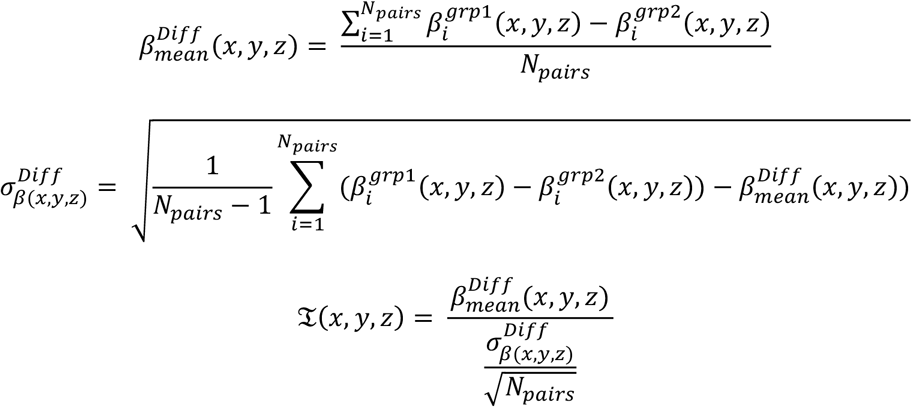

And for the region-time matrix,

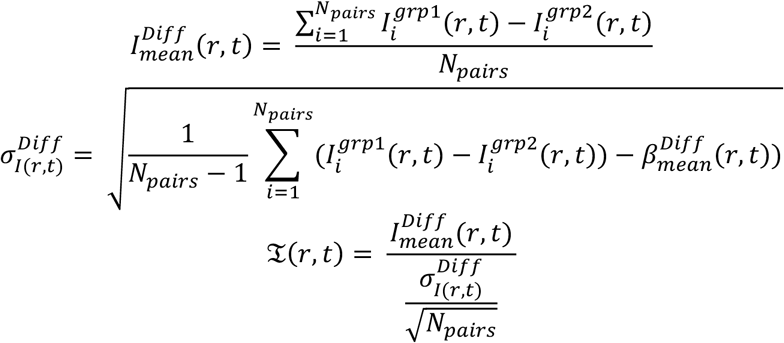

The one-sample and paired t-tests are implemented using the built-in MATLAB function *ttest*, while the two-sample t-test is performed using the function *ttest2*.

##### b.2. Visualization

Visualization serves as the culminating step in the data analysis process, arguably holding the utmost importance. It plays a pivotal role in enabling researchers to decipher and comprehend the extracted insights from the data. OfUSA provides a comprehensive collection of visualization tools to effectively represent both the spatial distribution and temporal dynamics of brain activity. For spatial distribution, multislice view and 3D rendering techniques are employed to pinpoint the localization of activity within the brain. Additionally, OfUSA offers region-time and time course views to visualize time course fluctuations, allowing researchers to track the evolution of brain activity over time. This comprehensive set of visualization tools empowers researchers to gain a deeper understanding of the complex patterns and relationships within the data.

###### b.2.1. Multislice view (mosaic view)

The multislice view, also known as the mosaic view, presents multiple slices in a single image. This visualization method enables researchers to effortlessly gain a comprehensive or partial overview of the fUS images (underlay images) or activation maps (overlay images). In OfUSA, researchers can select the specific slices to be displayed by specifying their indices in an array. For instance, the array [50, 70, 90, 95] would display the 50^th^, 70^th^, 90^th^, and 95^th^ slices.

The multislice view accommodates the display of various activation maps, including the signal change map for each trial [β_ABA (x, y, z)], the group mean signal change map [β_mean(x, y, z)], and the group t-value map [T(x, y, z)]. Consequently, the display range for the overlay data and threshold should be adjusted judiciously based on the characteristics of the activation map. For instance, the signal change map typically ranges between -15% and 15%, and a suitable threshold could be 3% (if the tSNR is 5 for each 36 trials, then the noise level of all 36 trials is approximately 3%). The t-value map theoretically ranges from -∞ to ∞, but in practice, it typically falls between -15 and 15. Appropriate thresholds for the t-value map could be T=2 (it is approximately the level of P<0.05 if there are 36 trials). Researchers should always adhere to the appropriate display range based on the loaded data.

When brain images are displayed in the coronal orientation, the top row or left column of the mosaic view corresponds to the anterior portion of the brain. Within each slice image, the right side represents the right hemisphere, while the left side represents the left hemisphere. The top and bottom edges correspond to the dorsal and ventral aspects of the brain, respectively.

When brain images are displayed in the sagittal orientation, the top row or left column of the mosaic view corresponds to the left portion of the brain. Within each slice image, the right side represents the anterior part of the brain, while the left side represents the posterior part. The top and bottom edges correspond to the dorsal and ventral aspects of the brain, respectively.

###### b.2.2. Region-time view (barcode view)

The region-time view, also known as the barcode view, seamlessly integrates spatial and temporal information into a single, informative figure. The arrangement of regions and time frames along the rows and columns, respectively, allows for a clear visualization of the time course for each region. This barcode-like representation effectively highlights activated regions and patterns of the hemodynamic response function.

The barcode view facilitates the visualization of various region-time matrix, including the single-trial matrix [I(r,t)], the group-averaged matrix [I_mean(r,t)], and the group t-value matrix [T(r,t)]. As discussed earlier, researchers should employ appropriate display ranges based on the characteristics of the region-time matrix in question. Signal change fluctuations typically fall within the range of -15% to 15%, while T-values invariably range from -15 to 15.

###### b.2.3. Time course view

The time course view empowers researchers to trace the hemodynamic response patterns of specific regions by plotting their time courses. The user-friendly interface enables researchers to manually select the ROI, allowing for a detailed examination of the hemodynamic response characteristics, including onset time, slew rate, peak value, and the activation tail.

Researchers can additionally visualize the standard deviation and average of the hemodynamic response function across trials, providing valuable insights into the reliability of the observed time course pattern. This visualization also serves as an effective tool for assessing the stability of baseline noise.

###### b.2.4. 3D Rendering

OfUSA additionally incorporates 3D rendering capabilities to assist researchers in identifying activation areas within a three-dimensional space. As outlined in the multislice view section, researchers should employ appropriate display ranges and thresholds to effectively visualize the 3D rendered surface. To prevent user overload from an excessive number of small activated clusters, only the top 50 clusters, ranked by cluster size, are displayed. VideoS1 illustrates an example of 3D rendering.

To render each cluster, OfUSA employs a stochastic algorithm to map activation strength onto the cluster’s surface. This algorithm assigns the highest activation value to the nearest surface point based on the Euclidean distance in 3D space. To expedite this process, we parallelize the surface mapping by randomly dividing the surface points into multiple subsets (each containing γ elements). For each surface subset, a randomly selected subset of activated voxels (with ω elements) is mapped to the nearest surface point. This iterative process continues until a majority of the activated voxels have been mapped. The stochastic algorithm can be summarized as follows:

> Step 1: Divide the surface into N_surf_ subsets, each containing γ elements. These subsets can be represented as Surface_i_(γ), where i ∈ N_surf_.
>
> Step 2: Repeat N_surf_ times: Randomly select a subset of voxels within the activated area, each with ω elements. These subsets can be denoted as Activated_j_(ω), where j ∈ N_act_.
>
> Step 3: Repeat N_act_ times: For each voxel in Activated_j_(ω), calculate the Euclidean distance between the voxel and the surface points Surface_i_(γ). If the voxel’s intensity exceeds the intensity of the nearest surface point, project the intensity onto the nearest ϵ points on the surface. The smoothing factor ϵ is set to 8 by default.
>
> Practically, we utilized MATLAB’s built-in functions to visualize the 3D rendered surface. The surface of the activated area was generated using the *isosurface* function, while the rendering was accomplished employing the *patch* function.

### B. Create data driven template

OfUSA creates a study-specific, data-driven template using an iterative alignment and averaging algorithm, which is illustrated in Figure S2. The automated registration at the core of this process utilizes the imregtform function in MATLAB. The procedure is as follows:

> Step 1: Initially, all anatomical images from each session (*IM*_*i*_, where i is the session number) are loaded.
>
> Step 2: An initial mean template, *Template*_0_, is computed by averaging these raw images.
>
> Step 3: The algorithm then enters an iterative loop. In each iteration k, every original image *IM*_*i*_ is realigned to the current template from the previous iteration *Template*_*k*−1_ to produce a set of newly registered images *IM*′_*i*_.
>
> Step 4: A new, refined template *Template*_*k*_ is then calculated by averaging these newly registered images *IM*′_*i*_.
>
> Step 5: This process of realignment and averaging (steps 3 and 4) is repeated 10 times. The output of the final iteration serves as the definitive data-driven template.

### C. Compare the preprocessing results

To compare the effect of different preprocessing steps, we used five indices to evaluate the quality of hemodynamic response, which includes the inverse of Noise level, SNR, CNR, D-prime, and the inverse of trial standard deviation.

The three indices can be presented in the following formula.

The SNR:

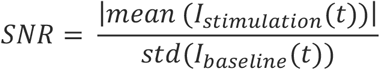

where *I*_*stimulation*_(*t*) is the hemodynamic signal intensity of the stimulation period.

The CNR:

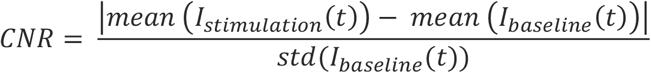

The D-prime (d’) :

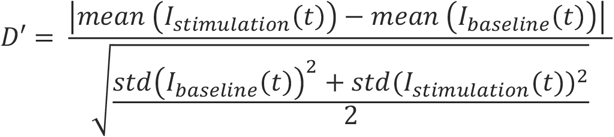

where *I*_*baseline*_(*t*) is the hemodynamic signal intensity of the baseline period, and s*td* denotes the standard deviation.

## Appendix II

### A. Mouse Experiment Setup

#### Animals and Preparation

A Wild-type C57BL/6J m mouse (∼25–30 g body weight; >8 weeks old; Janvier Labs) were used in this study. The fUS datasets were repurposed from previous fundamental research projects and were not generated specifically for the present work. All procedures were approved by the Animal Care Committee of Katholieke Universiteit Leuven, in compliance with national regulations on laboratory animal use and the European Union Directive 2010/63/EU on the protection of animals used for scientific purposes. To date, the approach has been applied exclusively to male mice, although it is not inherently limited to one sex. The mouse undergoing imaging eight times.

The animal preparation procedure adhered to the protocol outlined in our previous publication (Brunner et al., 2021). Briefly, the mice underwent cranial window surgery to expose the brain from bregma +3 mm to -7 mm. During the surgery, the mice were anesthetized with a ketamine/medetomidine mixture (0.1 mL/20 g), and their body temperature was maintained at 37 degrees Celsius using an in-house heating system. To minimize inflammation and pain, the animals received post-surgical treatment with analgesic (buprenorphine 0.1 mg/kg), antibiotic (cefazoline 300 mg/kg), and anti-inflammatory (dexamethasone 0.5 mg/kg). Following a five-day recovery period, the mice underwent gradual habituation to the head-fixed position.

#### Experimental Paradigm

This experiment employed visual and tactile stimuli. Visual stimulation consisted of a checkerboard pattern presented on the right side of the LCD screen positioned 20 cm from the subject. Tactile stimulation was delivered to the entire right whisker pad using a 3D-printed comb driven by a piezoelectric actuator (PiezoDrive BA6020). The maximum deflection of the comb is 0.7 cm. All stimuli lasted for 5 seconds. Figure 4 illustrates the experimental setup and the stimuli.

Each trial comprised a 10-second baseline period, followed by a 5-second stimulation period, and concluded with a 15-second recovery period, resulting in a total trial duration of 30 seconds. Three experimental conditions were presented in each session: visual [V] and whisker [W]. Each condition was randomly presented 20 times per session. A high-quality anatomy image was obtained after the experiment for registration purposes. Each session lasted approximately one hour and ten minutes.

#### vfUS imaging

For acquiring the vfUS imaging, we employed the acquisition toolbox with the following hardware setup (Brunner et al., 2020, 2021). In summary, we utilized a 2D-array transducer (MAT 15.0/32 x 32, Vermon, France) with a surface area of 9.6 x 9.6 mm, positioned on the cranial windows prepared earlier. We employed a central excitation frequency of 15 MHz with a bandwidth of 14 MHz (encompassing 8 MHz to 22 MHz). The sampling rate for the scattered echoes was 60 MHz. The elements on the probe were divided into four sectors, each containing 8 x 32 elements. 3D compound plane waves were generated by transmitting and receiving from these sectors. Ultimately, we reconstructed vfUS whole-brain images with a field of view (FOV) of 9.6 x 8.1 x 7.0 mm in the left-right, anterior-posterior, and dorsal-ventral directions, respectively. The corresponding voxel size was 0.15 x 0.15 x 0.1 mm in the left-right, anterior-posterior, and dorsal-ventral directions, respectively. Each brain volume had a frame rate of 2 Hz.

### B. Primate Experiment Setup

#### Animals and Preparation

A single male rhesus macaque (Macaca mulatta; 9.5 kg, 9 years old) was utilized for this study. All housing and procedures were conducted at the KU Leuven Medical School primate facility. The animal was socially housed in a group enclosure with access to physical and mental enrichment (e.g., toys, music) and was exposed to a 12-hour natural and artificial light-dark cycle. The subject’s health was continuously monitored by trained technical staff, veterinary staff, and experimenters.

The surgical procedures were performed as previously described in (Vanduffel et al., 2001). Briefly, an MR-compatible headpost and recording chamber were surgically implanted. The implant was secured using dental cement, and the recording chamber was positioned according to pre-operative anatomical images to ensure access to areas of the dorsal and ventral visual streams. All surgical and experimental protocols were in agreement with institutional (KU Leuven Medical School: Ethische Commissie Dierproeven), national, and European guidelines (Directive 2010/63/EU).

#### Experimental Paradigm

The subject was trained via operant conditioning with a liquid juice reward to perform a passive fixation task. During all experimental sessions, the subject was head-fixed and seated in the sphinx position. To minimize motion artifacts, the subject was also trained to position its hands in a box located directly in front of and below its head.

Visual stimuli were presented on a 32-inch screen (3840 x 2160 pixels) positioned 40 cm from the subject, while eye position was tracked at 120 Hz using an infrared camera system (ISCAN; Woburn, MA, USA) focused on the right eye. A trial was initiated when the subject maintained fixation on a central white dot within a 2x2-degree window. Following a variable inter-trial interval (ITI), a circular, colorful checkerboard stimulus with a radius of 40 visual degrees was presented centrally for 8 seconds, during which it flashed at 10 Hz. The subject received a reward on a fixed schedule for maintaining fixation throughout the stimulus presentation. Each experimental run lasted approximately 18 minutes, comprised approximately 18 trials, and utilized an ITI that varied between 18 and 22 seconds in 0.5-second increments. The probability of each ITI was weighted in a descending linear fashion, such that the 18-second ITI was nine times more likely to occur than the 22-second ITI.

#### vfUS imaging

The vfUS imaging acquisition was adapted from the technique described in (Brunner et al., 2021). We utilized the same 2D-array transducer (MAT 15.0/32 x 32, Vermon, France) as in the prior rodent study. This probe has a central excitation frequency of 15 MHz, which permits a penetration depth of approximately 1 cm into the brain. Ultimately, we reconstructed vfUS images with a field of view (FOV) of 9.6 x 8.1 x 100 mm in the left-right, anterior-posterior, and dorsal-ventral directions, respectively. The corresponding voxel size was 0.15 x 0.15 x 0.1 mm in the left-right, anterior-posterior, and dorsal-ventral directions, respectively. Each brain volume was acquired at a frame rate of 1.4 Hz.

