## supplementary for "OfUSA: OpenfUS Analyzer, a versatile open-source framework for the analysis and visualization of functional ultrasound imaging data across animal models"

#### Figure S1

a. The pipeline for individual analysis

| Process Name | Status |
|---|---|
| Loop Pipeline (1) | Ready |
| Set Parameters (2) | Wait |
| Regressor (3) | Wait |
| ---fill the parameters above--- | |
| ---please do not change parameters below--- | |
| ---Preprocess--- | |
| Quality Control (4) | Wait |
| Quality Control (5) | Wait |
| Temporal Filter (6) | Wait |
| Temporal Filter (7) | Wait |
| ---Timecourse--- | |
| Segmentation ABA (8) | Wait |
| Segmentation ABA (9) | Wait |
| Average across data (10) | Wait |
| Average across data (11) | Wait |
| ---ActivationMap--- | |
| GLM Analysis (12) | Wait |
| GLM Analysis (13) | Wait |
| Average across data (14) | Wait |
| Average across data (15) | Wait |
| Registration: ABA (16) | Wait |
| Registration: ABA (17) | Wait |

b. The pipeline for group analysis

| Process Name | Status |
|---|---|
| ---Select time course data--- | |
| Select Data (1) | Ready |
| ---Select activation maps--- | |
| Select Data (2) | Wait |
| Set Parameters (3) | Wait |
| ---please fill the parameters above--- | |
| ---please do not change the parameter below--- | |
| ---Timecourse--- | |
| Group T-test (4) | Wait |
| Group T-test (5) | Wait |
| Timecourse View (6) | Wait |
| Barcode View (7) | Wait |
| ---BrainMap--- | |
| Group T-test (8) | Wait |
| Group T-test (9) | Wait |
| Mosaic View (10) | Wait |
| Regions Report (11) | Wait |

c. Interface for registering data

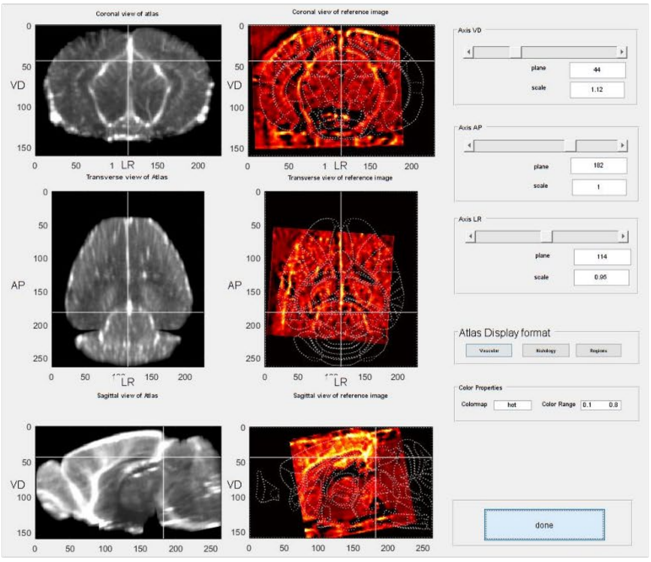


d. Interface for selecting data for a batch process

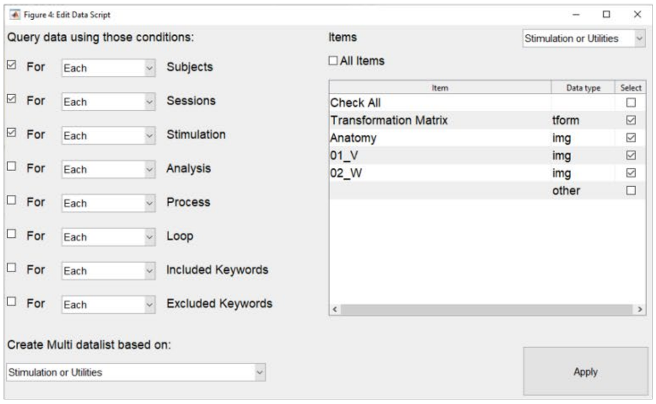


e. Interface for defining experimental regressor

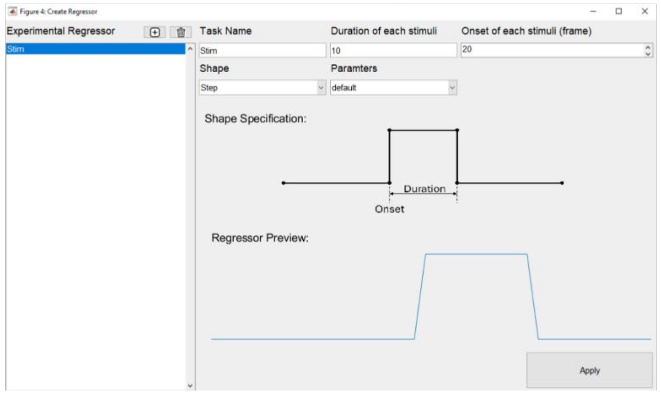


Figure S1. Examples of analysis pipelines and interfaces for the rodent study.
Panel (a) shows the individual-level analysis pipeline, consisting of 17 processes. Panel (b) presents the group-level pipeline, composed of 11 processes. Panel (c) displays the manual registration interface, with the reference vascular template (left), the subject's high-resolution anatomical image (middle), and slice adjustment parameters (right). Panel (d) illustrates the interface for batch data selection, where query parameters are listed on the left and corresponding values on the right. Panel (e) depicts the interface for defining the experimental regressor, enabling specification of stimulus shape, onset time, and duration.

### Figure S2

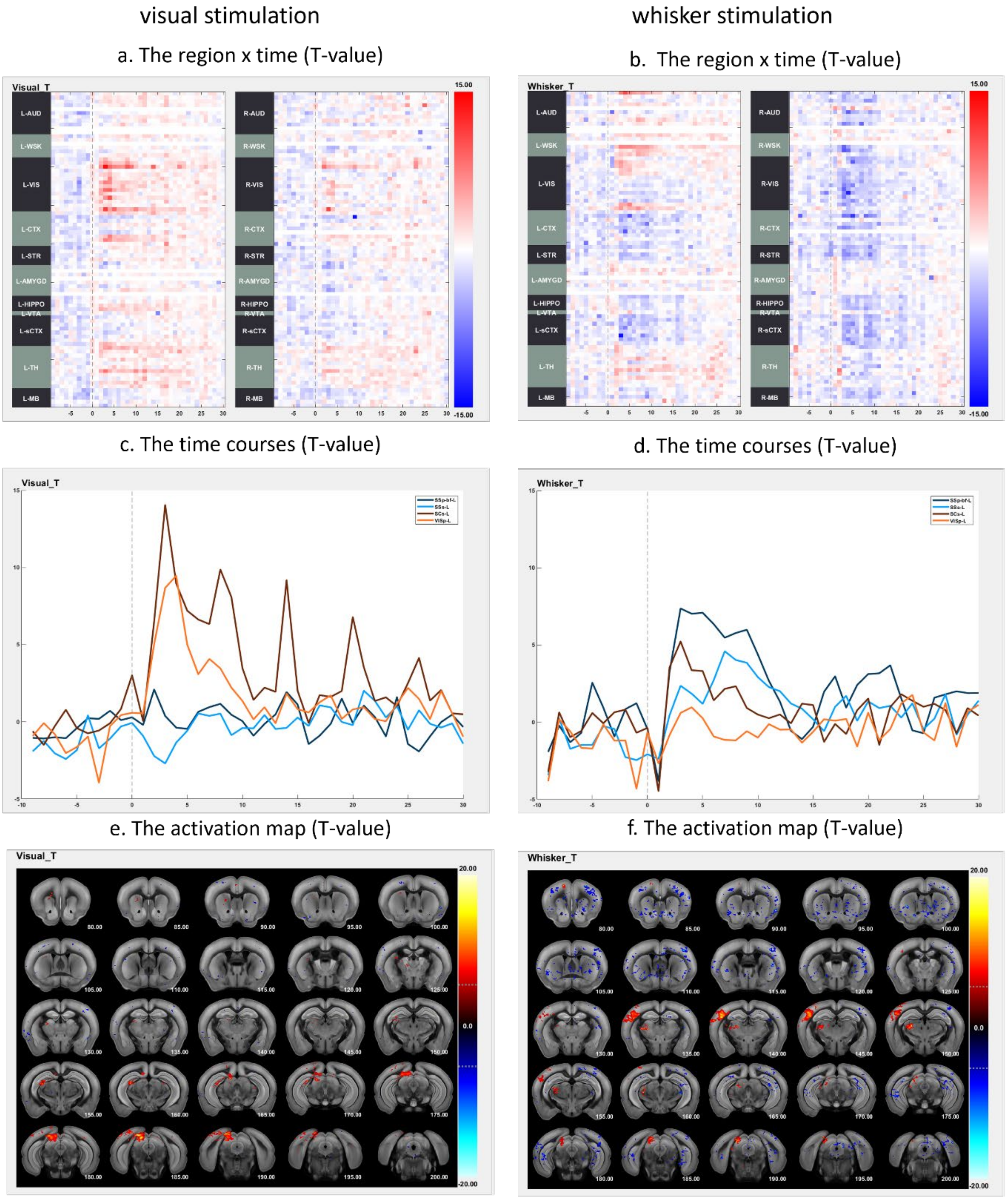


Figure S2. Typical results from the rodent study.

Panels (a, b) present region-by-time plots of time courses across multiple brain regions. Panels (c, d) display representative time courses for visual regions (VISp-L, SCs-L, orange) and somatosensory regions (SSp-bfd-L, SSs-L, blue). Panels (e, f) show the group-level activation maps. Reported activations are statistically significant according to a one-sample t-test ($T > 5.4$, $p < 0.001$ uncorrected, $df = 7$, two-tailed).

### Figure S3

a. Workflow to create data driven template

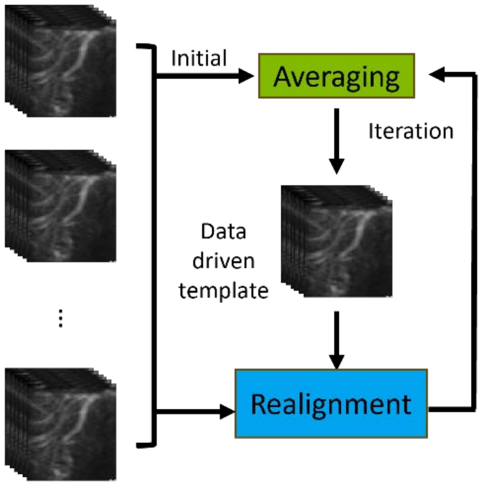


b. One of images in Monkey visual area

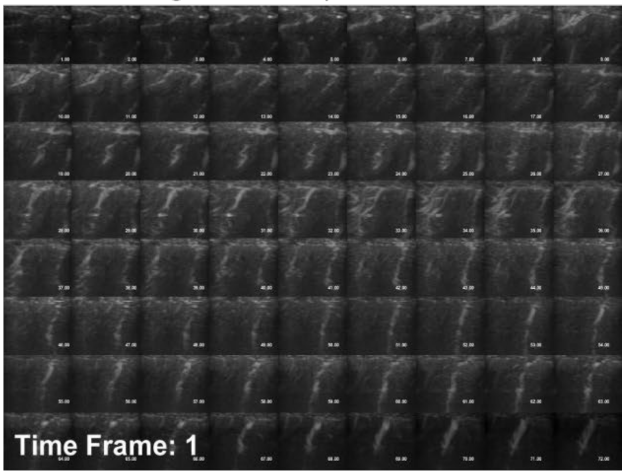


c. The data driven template ( from 8 sessions)

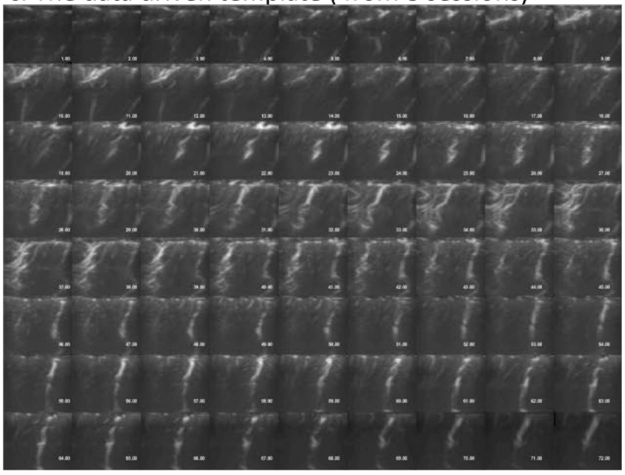

d. The data driven template with labeled ROI

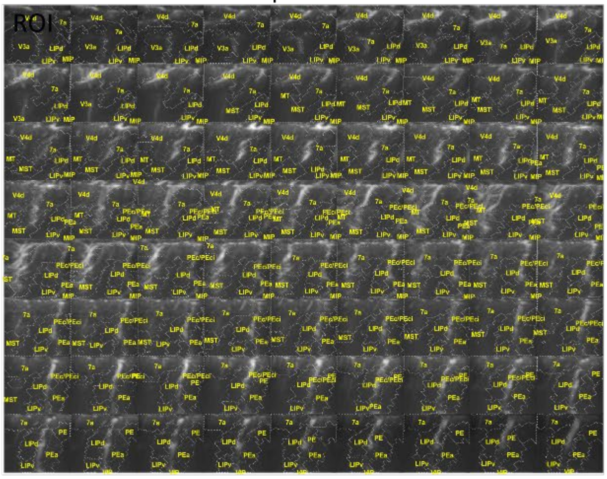

e. The pipeline for individual analysis

| Process Name | Status |
|---|---|
| Loop Pipeline (1) | Ready |
| ---select template--- | |
| Select Data (2) | Wait |
| ---Fill in the parameters above--- | |
| ---Please do not modify the parameters below--- | |
| ---Preprocess--- | |
| Quality Control (3) | Wait |
| Temporal Filter (4) | Wait |
| Realignment (5) | Wait |
| ---Analysis--- | |
| Extract Timecourse (6) | Wait |
| Average across data (7) | Wait |
| GLM Analysis (8) | Wait |

f. The pipeline for group analysis

| Process Name | Status |
|---|---|
| ---TemporalDataSelection--- | |
| Select Data (1) | Ready |
| ---BrainMapDataSelection--- | |
| Select Data (2) | Wait |
| ---Template_Selection--- | |
| Select Data (3) | Wait |
| ---Set Parameters--- | |
| Set Parameters (4) | Wait |
| ---Please edit the processes above--- | |
| ---Please do not edit the following processes--- | |
| ---Timecourse--- | |
| Average across data (5) | Wait |
| Barcode View (6) | Wait |
| Timecourse View (7) | Wait |
| ---BrainMap--- | |
| Fixed Effect Analysis (8) | Wait |
| Mosaic View (9) | Wait |
| Regions Report (10) | Wait |

Figure S3. Workflow and pipelines of the data-driven template for the primate study.

Panel (a) shows the iterative workflow for generating a data-driven template from primate data. All session images are averaged to create an initial template, after which raw images are realigned to this template and re-averaged. This realignment and averaging procedure is repeated for 10 iterations to obtain the final template. Panel (b) presents an example of a primate anatomical image prior to realignment. Panel (c) displays the final data-driven template generated through the iterative workflow, and panel (d) shows the final template with labeled regions of interest. Panel (e) illustrates the individual-level analysis pipeline, consisting of 8 processes, and panel (f) presents the group-level pipeline, composed of 10 processes.

Figure S4

a. Activation map: T-value

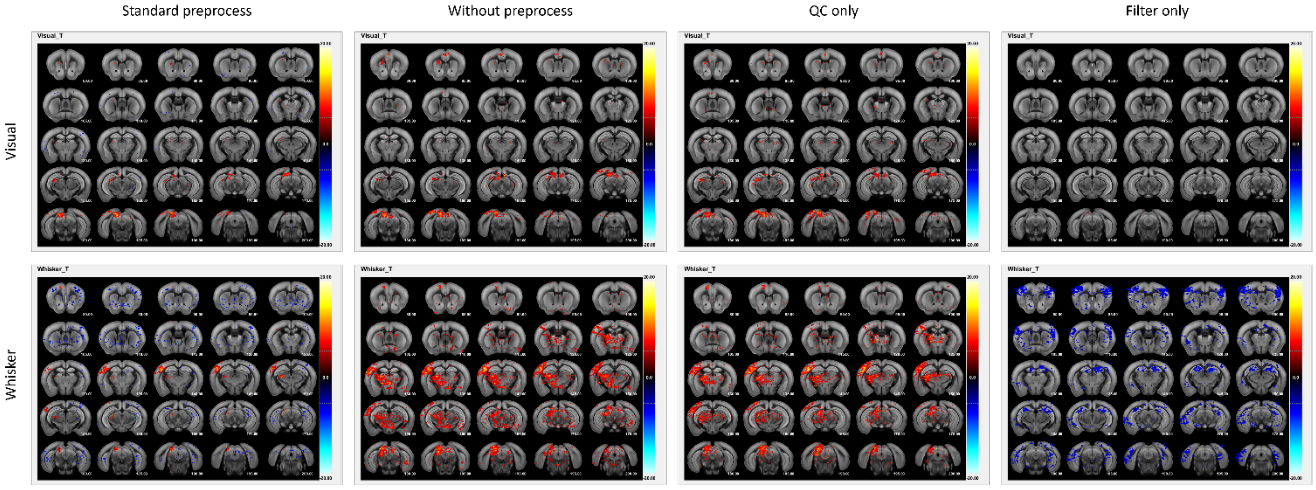


b. Region x Time courses : signal change

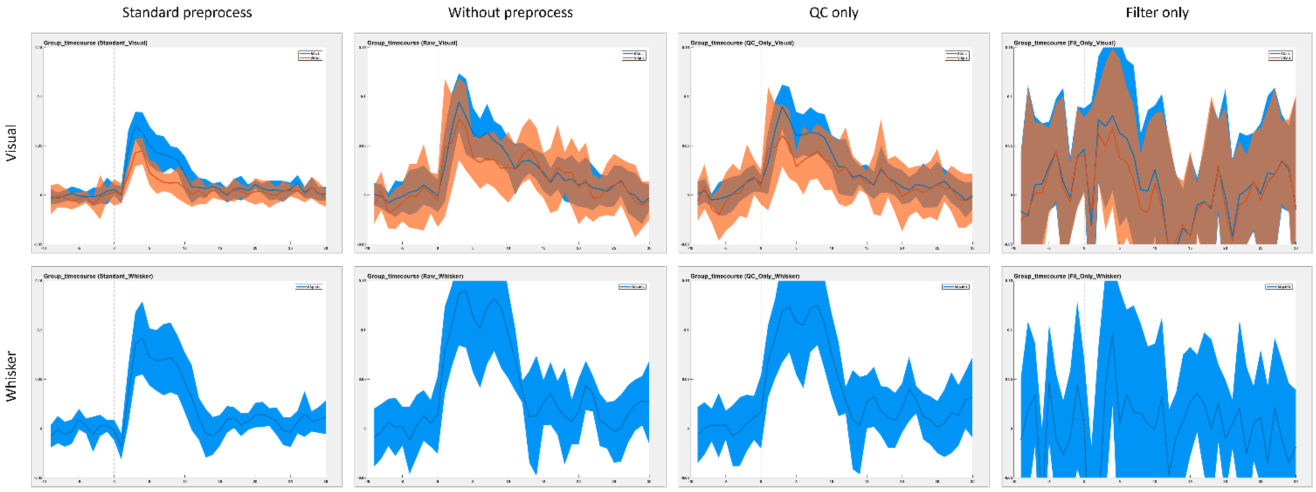


c. Comparison

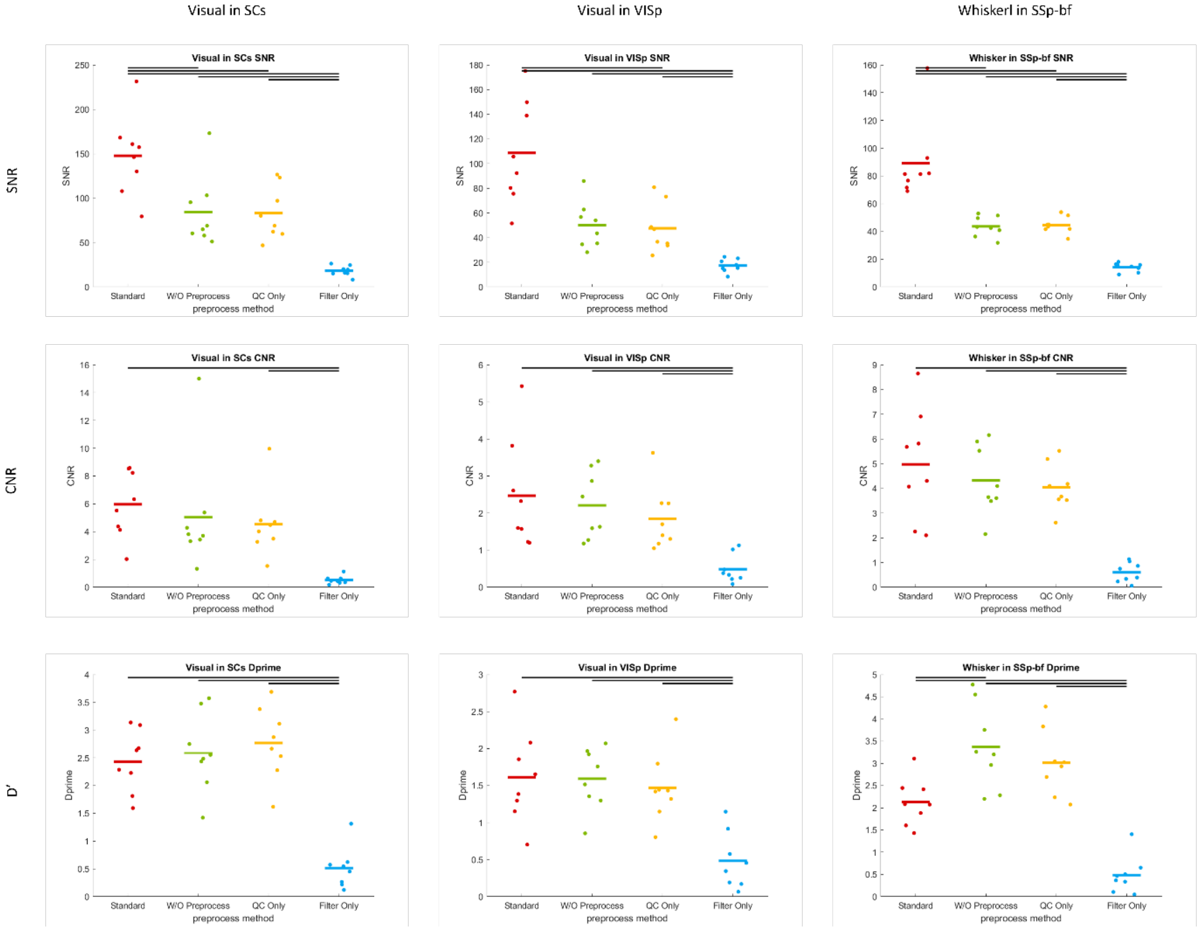

Figure S4. Comparison of preprocessing methods in the rodent study.

Panel (a) shows group-level activation maps for visual and whisker stimulation obtained under four preprocessing conditions: standard (quality check and filtering), no preprocessing, quality check (QC) only, and filtering only. Panel (b) presents the corresponding time courses from selected regions of interest (VISp and SCs for visual stimulation; SSp-bfd for whisker stimulation) for each method. Panel (c) provides a quantitative comparison of the time courses from panel (b) using signal-to-noise ratio (SNR), contrast-to-noise ratio (CNR), and d-prime (d′). Significant differences between methods are indicated by bars, based on paired t-tests ($p < 0.05$, corrected for multiple comparisons).

Figure S5

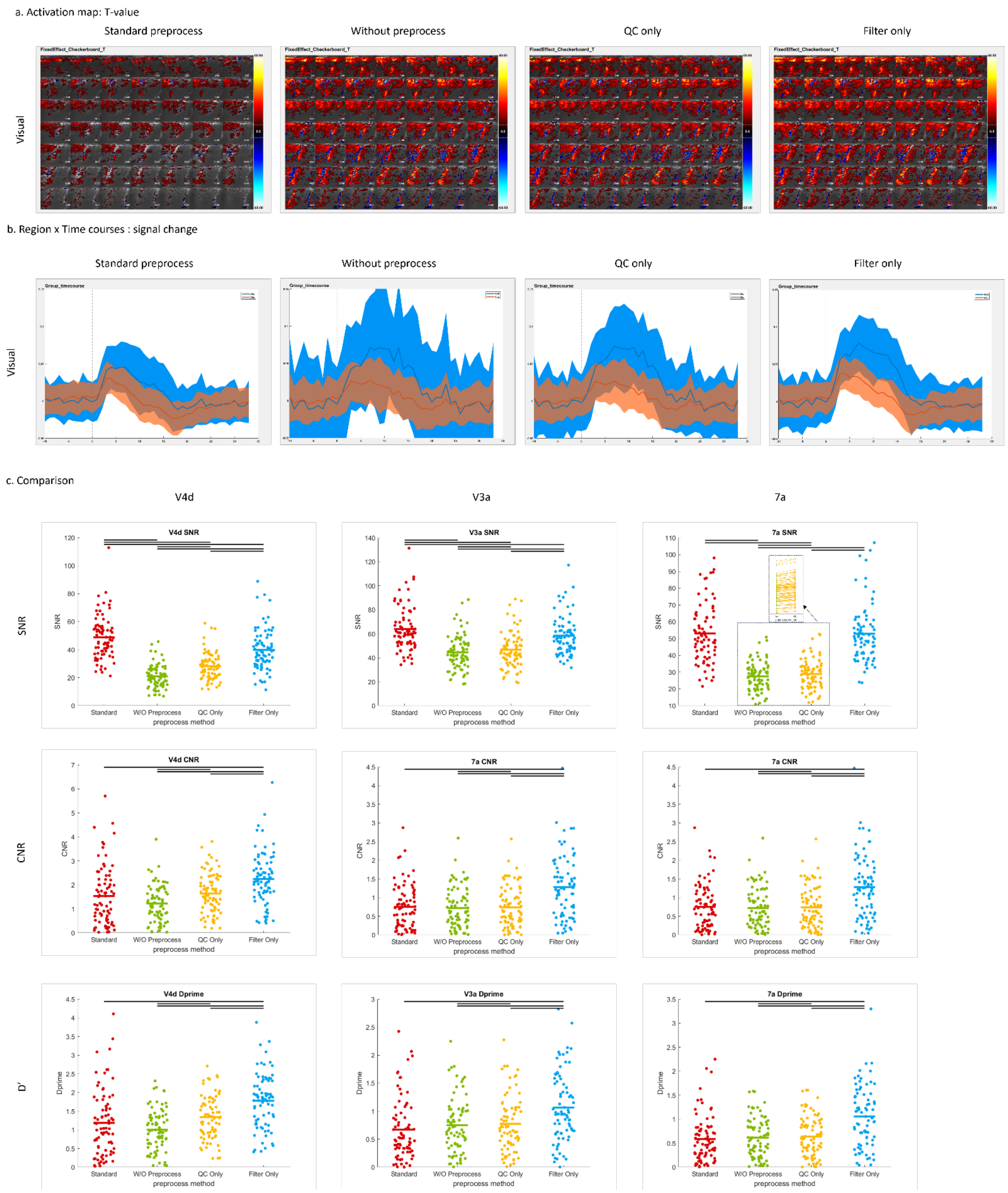


Figure S5. Comparison of preprocessing methods in the primate study.

Panel (a) shows group-level activation maps for checkerboard stimulation under four preprocessing conditions: standard (quality check and filtering), no preprocessing, quality check (QC) only, and filtering only. Panel (b) presents the corresponding time courses from selected regions of interest (V4d and V3a) for each method. Panel (c) provides a quantitative comparison of the time courses from V4d, V3a, and 7a using signal-to-noise ratio (SNR), contrast-to-noise ratio (CNR), and d-prime (d′). Significant differences between methods are indicated by bars, based on paired t-tests ($p < 0.05$, corrected for multiple comparisons).

**Videos**

Video S1. Three-dimensional rendering of activation during visual stimulation in the rodent study.

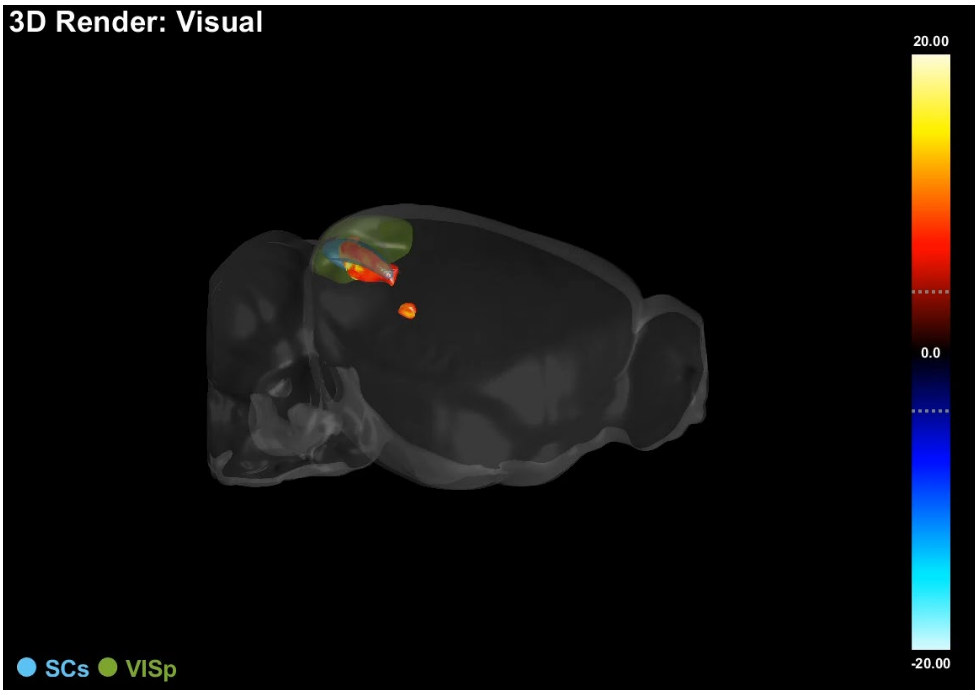


https://zenodo.org/records/17132116/files/3DRender_Visual.mp4

Video S2. Three-dimensional rendering of activation during whisker stimulation in the rodent study.

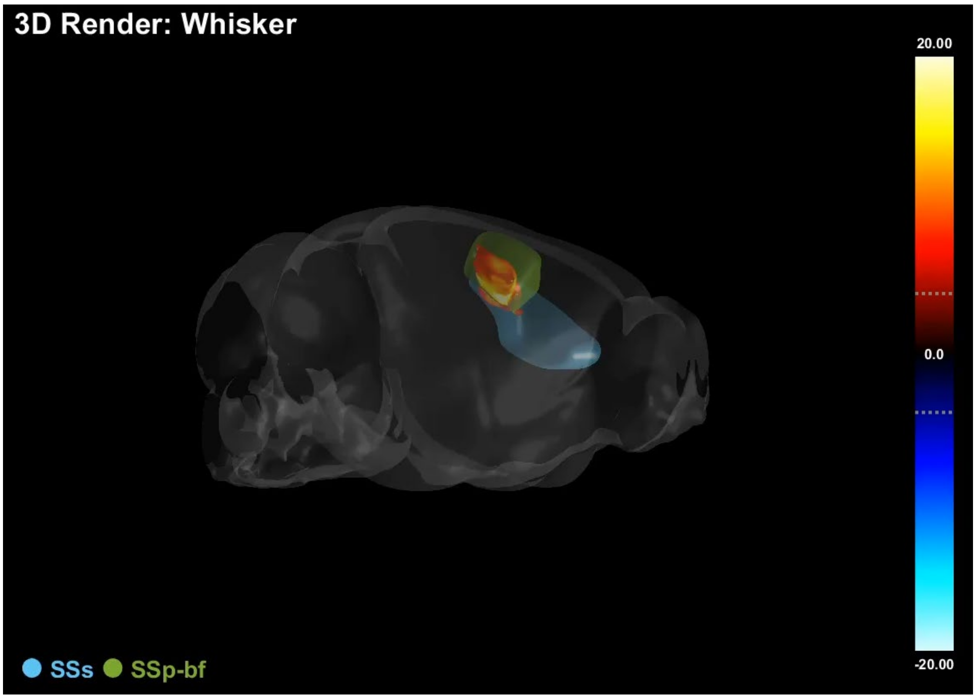


https://zenodo.org/records/17132116/files/3DRender_Whisker.mp4

Table S1. Metadata stored in the JSON file for data used in OfUSA.

| Parameters Name | Function |
|---|---|
| **File_Path** | The JSON file path. |
| **Data_Path** | The corresponding NIFTI file path. |
| **SubjectID** | The Mouse ID. |
| **SessionID** | The ID of the session. |
| **StimulusID** | The stimulus ID is the name of the stimulus or experimental condition. |
| **TrialID** | The Trial ID. |
| **AnalysisID** | The Analysis ID |
| **ProcID** | The process ID |
| **DataCategory** | Description of the data type, it could be raw data (raw), analyzed data (ana), utilities (utl). |
| **Orientation** | Description of the data orientation, it could be coronal (cor), sagittal (sag) or transverse (trans). |
| **Dimension** | The 3 values array denotes the the image dimension in Left-Right, Anterior-Posterior, Ventral-Dorsal directions, respectively. |
| **Voxel_Size** | The 3 values array denotes the voxel size of image in Left-Right, Anterior-Posterior, Ventral-Dorsal directions, respectively. The unit is micrometer. |
| **Repetition** | The repetition denotes the number of time points. |
| **SampleInterval** | The sampling interval is the inverse of the sampling rate. The unit is second. |

Table S2. List of ROIs associated with sensory areas in the mouse.

| ROI Name | Full Name | Cluster Name | Cluster Full Name |
|---|---|---|---|
| **AUDd** | Dorsal auditory area | AUD | Auditory-related area |
| **AUDp** | Primary auditory area | AUD | Auditory-related area |
| **AUDpo** | Posterior auditory area | AUD | Auditory-related area |
| **AUDv** | Ventral auditory area | AUD | Auditory-related area |
| **MOs** | Secondary motor area | AUD | Auditory-related area |
| **MGN** | Medial geniculate complex | AUD | Auditory-related area |
| **IC** | Inferior colliculus | AUD | Auditory-related area |
| **LL** | Lateral lemniscus | AUD | Auditory-related area |
| **SOC** | Superior olivary complex | AUD | Auditory-related area |
| **TRN** | Tegmental reticular nucleus | AUD | Auditory-related area |
| **CN** | Cochlear nucleus | AUD | Auditory-related area |
| **SSp-bf** | Primary somatosensory area, barrel field | WSK | Whisker-related area |
| **SSs** | Supplemental somatosensory area | WSK | Whisker-related area |
| **PO** | Posterior complex of the thalamus | WSK | Whisker-related area |
| **VP** | ventral posterior complex of the thalamus | WSK | Whisker-related area |
| **PSV** | Principal sensory nucleus of the trigeminal | WSK | Whisker-related area |
| **SPV** | Spinal nucleus of the trigeminal | WSK | Whisker-related area |
| **VISa** | Anterior visual area | VIS | Visual-related area |
| **VISal** | Anterolateral visual area | VIS | Visual-related area |
| **VISam** | Anteromedial visual area | VIS | Visual-related area |
| **VISl** | Lateral visual area | VIS | Visual-related area |
| **VISli** | Laterointermediate visual area | VIS | Visual-related area |
| **VISp** | Primary visual area | VIS | Visual-related area |
| **VISpl** | Posterolateral visual area | VIS | Visual-related area |
| **VISpm** | Posteromedial visual area | VIS | Visual-related area |
| **VISpor** | Postrhinal area | VIS | Visual-related area |
| **VISrl** | Rostrolateral visual area | VIS | Visual-related area |
| **dLG** | Dorsal part of the lateral geniculate complex | VIS | Visual-related area |
| **SCd** | Superior colliculus, deep layers | VIS | Visual-related area |
| **SCi** | Superior colliculus, intermediate layers | VIS | Visual-related area |
| **SCs** | Superior colliculus, superficial layers | VIS | Visual-related area |
| **ACA** | Anterior cingulate area | CTX | Cortex |
| **AIa** | Agranular insular area | CTX | Cortex |
| **ECT** | Ectorhinal | CTX | Cortex |
| **ILA** | Infralimbic area | CTX | Cortex |
| **MOp** | Primary motor area | CTX | Cortex |
| **PERI** | Perirhinal area | CTX | Cortex |
| **PL** | Prelimbic area | CTX | Cortex |
| **RSP** | Restrosplenial area | CTX | Cortex |
| **TEa** | Temporal association areas | CTX | Cortex |

| | | | |
|---|---|---|---|
| **sAMY** | Amygdalar area | STR | Striatum |
| **Cpa** | Anterior caudoputamen | STR | Striatum |
| **CPm** | Medial caudoputamen | STR | Striatum |
| **CPc** | Caudal caudoputamen | STR | Striatum |
| **LS** | Lateral septal nucleus | STR | Striatum |
| **PAA** | Piriform-amygdalar area | AMYGD | amygdalar |
| **MEA** | Medial amygdalar nucleus | AMYGD | amygdalar |
| **LA** | Lateral amygdalar nucleus | AMYGD | amygdalar |
| **COA** | Cortical amygdalar area | AMYGD | amygdalar |
| **BLAd** | Basolateral amygdalar nucleus, dorsal part | AMYGD | amygdalar |
| **BLAv** | Basolateral amygdalar nucleus, ventral part | AMYGD | amygdalar |
| **BMA** | Basomedial amygdalar nucleus | AMYGD | amygdalar |
| **HATA** | Hippocampo-amygdalar transition area | AMYGD | amygdalar |
| **CA1** | CA1 subfield | HIPPO | Hippocampus |
| **CA2** | CA2 subfield | HIPPO | Hippocampus |
| **CA3** | CA3 subfield | HIPPO | Hippocampus |
| **DG** | Dentate gyrus | HIPPO | Hippocampus |
| **VTA** | Ventral tegmental area | VTA | Ventral tegmental area |
| **ENT** | Entorhinal area | sCTX | subcortical area |
| **SUB** | Subiculum | sCTX | subcortical area |
| **ACB** | Nucleus accumbens | sCTX | subcortical area |
| **BST** | Bed nuclei of the stria terminalis | sCTX | subcortical area |
| **CLA** | Claustrum | sCTX | subcortical area |
| **FS** | Fundus of striatum | sCTX | subcortical area |
| **GP** | Globus pallidus | sCTX | subcortical area |
| **MS** | Medial septal nucleus | sCTX | subcortical area |
| **ATN** | Anterior group of dorsal nucleus | TH | Thalamus |
| **CL** | Central lateral nucleus of the thalamus | TH | Thalamus |
| **CM** | Central medial nucleus of the thalamus | TH | Thalamus |
| **IGL** | Intergeniculate leaflet of the lateral geniculate complex | TH | Thalamus |
| **IMD** | Intermediodorsal nucleus of the thalamus | TH | Thalamus |
| **LGv** | Ventral part of the lateral geniculate complex | TH | Thalamus |
| **LP Eth** | Lateral posterior nucleus of the thalamus | TH | Thalamus |
| **MD** | Mediodorsal nucleus of thalamus | TH | Thalamus |
| **PPnT** | Posterior Paralaminar nuclei of the thalamus | TH | Thalamus |
| **VAL** | Ventral anterior-lateral complex of the thalamus | TH | Thalamus |
| **VM** | Ventral medial nucleus of the thalamus | TH | Thalamus |
| **CUN** | Cuneiform nucleus | MB | Midbrain |
| **MRN** | Midbrain reticular nucleus | MB | Midbrain |
| **PAG** | Periaqueductal gray | MB | Midbrain |
| **PBG** | Parabigeminal nucleus | MB | Midbrain |
| **SN** | Substantia nigra | MB | Midbrain |

Table S3. Behavioral metadata stored in the JSON file of the behavioral regressor used in OfUSA.

| Parameters Name | Function |
|---|---|
| **Sti_Name** | The name of the stimulation. If there is more than one kind of stimuli, use a comma to separate them. |
| **Sti_Duration** | The duration of the stimuli is measured in frames. If there is more than one kind of stimulus, use a comma to separate them. |
| **Sti_Onset** | If there are multiple kinds of stimuli, their onset frames should be provided as a comma-separated sequence of bracketed lists. Each set of square brackets [] represents a distinct kind of stimulus.<br><br>Format: [kind1_onset1, kind1_onset2, ...], [kind2_onset1, kind2_onset2, ...]<br><br>Within each bracketed list, individual onset times (in frames) are separated by commas. The order of these bracketed lists should align with the defined order of your stimulus kinds. |
| **Sti_Shape** | This parameter defines the temporal profile of the stimulation's intensity: use "step" for a block design where intensity is constant throughout its duration; use "Triangular" for a shape that gradually increases to a peak and then gradually decreases; and use "Trapezoidal" for a shape that gradually increases, holds at a plateau of peak intensity, and then gradually decreases. If there is more than one kind of stimulus, use a comma to separate them. |
| **Sti_Par** | This extra parameters provide additional options for defining the shape of the stimulation. |
| **Reg_num** | The total number of stimulus kinds. |

Table S4. List of ROIs in the monkey study.

| ROI Name | Full Name | Cluster Name |
|---|---|---|
| **PEc/PEci** | Caudal and Cingulate portion of area PE | Parietal |
| **PEa** | Anterior part area of area PE | Parietal |
| **MIP** | Medial Intraparietal Area | Intraparietal |
| **LIPd** | Dorsal Lateral Intraparietal Area | Intraparietal |
| **LIPv** | Ventral Lateral Intraparietal Area | Intraparietal |
| **MST** | Medial Superior Temporal Area | Temporal |
| **7a** | Area 7a | Parietal |
| **MT** | Middle Temporal Area | Temporal |
| **V4d** | Dorsal Visual Area 4 | Visual |
| **V3a** | Visual Area 3A | Visual |
